# Transneuronal Dpr12/DIP-δ interactions facilitate compartmentalized dopaminergic innervation of *Drosophila* mushroom body axons

**DOI:** 10.1101/834515

**Authors:** Bavat Bornstein, Idan Alyagor, Victoria Berkun, Hagar Meltzer, Fabienne Reh, Hadas Keren-Shaul, Eyal David, Thomas Riemensperger, Oren Schuldiner

## Abstract

The mechanisms controlling wiring of neuronal networks are largely unknown. The stereotypic architecture of the *Drosophila* mushroom-body (MB) offers a unique system to study circuit assembly. The adult medial MB γ-lobe is comprised of a long bundle of axons that wires with specific modulatory and output neurons in a tiled manner defining five distinct zones. We found that the immunoglobulin superfamily protein Dpr12 is cell-autonomously required in γ-neurons for their developmental regrowth into the distal γ4/5 zones, where both Dpr12 and its interacting protein, DIP-δ, are enriched. DIP-δ functions in a subset of dopaminergic neurons that wire with γ-neurons within the γ4/5 zone. During metamorphosis, these dopaminergic projections arrive to the γ4/5 zone prior to γ-axons, suggesting that γ-axons extend through a prepatterned region. Thus, Dpr12/DIP-δ transneuronal interaction is required for γ4/5 zone formation. Our study sheds light onto molecular and cellular mechanisms underlying circuit formation within subcellular resolution.

## Introduction

The precise connectivity between neurons is crucial for the function of neural circuits in vertebrates and invertebrates. The formation of neural circuits is especially complex as it is a multi-step process that involves guidance of axons and dendrites belonging to distinct neurons, as well as the identification of subcellular zones on the target cell onto which synapses are formed. Despite its fundamental nature, the molecular and cellular mechanisms underlying development of neural circuits remain mostly poorly understood.

Given its unique development, connectivity and function, the *Drosophila* mushroom body (MB), offers an attractive model to study the mechanisms of neuronal circuit formation and maturation. The adult MB, implicated in associated learning (Fiala, 2007; Gerber et al., 2004; Heisenberg, 2003; Modi et al., 2020; Owald and Waddell, 2015), is comprised of intrinsic as well as extrinsic neurons (Aso et al., 2014a; Tanaka et al., 2008). Intrinsic MB neurons are derived from four identical neuroblasts which sequentially give rise to three major classes of unipolar neurons: γ, α’/β’ and α/β, which are collectively known as Kenyon Cells (KCs). Axons from each KC type bundle together to form five MB lobes in the adult brain - the vertical α and α’ lobes and the medial γ, β and β’ lobes (Figure 1A, Crittenden et al., 1998). KCs form well-defined circuits with MB extrinsic neurons, which include MB output neurons (MBONs) and modulatory neurons, mostly dopaminergic (DANs). The processes of MBONs and DANs innervate the MB lobes at distinct and stereotypic locations thereby forming discrete zones, which are also known as compartments (Due to a potential confusion between cell intrinsic compartments such as the axon initial segment, here we use the term zone to describe these lobe-compartments; Aso et al., 2014a; Tanaka et al., 2008). For example, the γ-lobe, which is comprised of γ-KC axons, is innervated by extrinsic neurons in five distinct axonal zones termed γ1-γ5. Thus, each γ-zone is defined by stereotypic inputs from specific DANs and MBONs (Figure 1A). Each γ-axon extends throughout the entire lobe and forms synaptic boutons with different partners within each zone. Remarkably, a recent study has shown that boutons within the same KC, but in different zones, often exhibit distinct calcium dynamics (Bilz et al., 2020). Finally, these zones have distinct functional roles. DANs innervating the γ1-γ2 zones are associated with aversive memory, while DANs innervating the γ4-5 zones promote appetitive memory (Aso et al., 2014a; Cognigni et al., 2018; Cohn et al., 2015). Despite the functional importance, the cellular and molecular mechanisms that control MB circuit and zone formation are not known.

**Figure 1.**
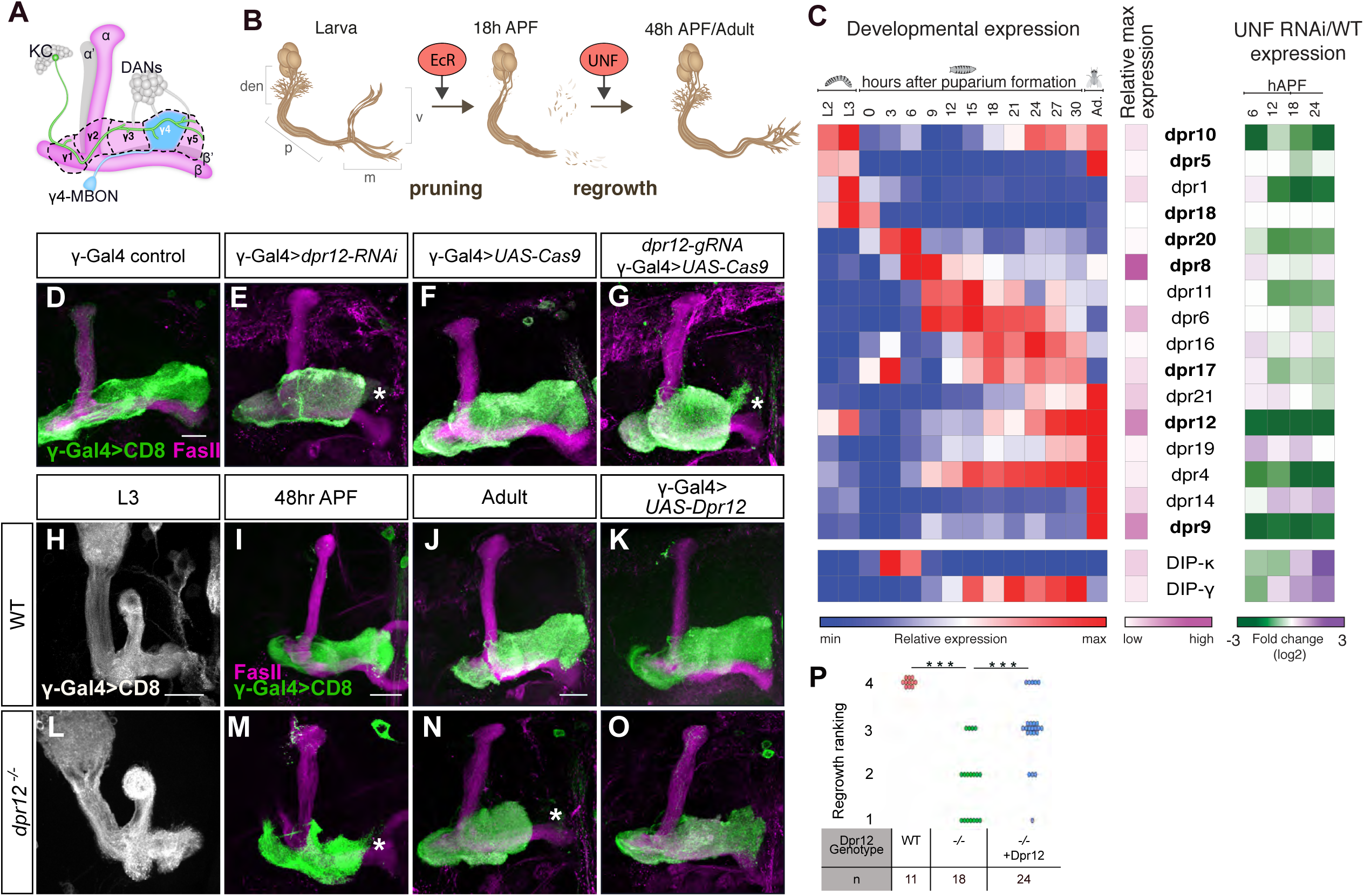
Dpr12 is required for full extension of γ-KCs. (A) Schematic representation of the adult MB. The bundled γ-axons (an example of which is depicted in green) form the lobe. Stereotyped and tiled innervation of the lobe by Dopaminergic (DANs, gray) and MB output neurons (MBONs, an example of a γ4 MBON is shown in cyan) define the γ1-γ5 zones. (B) Schematic representation of neuron remodeling of γ-KCs and its regulation by the nuclear receptors EcR and UNF. den: dendrites; p: axon peduncle; m/v: medial and vertical lobes. (C) Dynamic expression of Dpr and DIPs during γ-KCs development. Left: Heatmap depicting the relative expression patterns of Dprs and DIPs in γ-KCs during development. Middle: Magenta intensity depicts the peak expression of each gene during development relative to other Dprs and DIPs. Right: Expression change of Dprs and DIPs while knocking down the UNF transcription factor compared to WT γ-KCs. Dprs highlighted in bold were tested in the RNAi mini-screen (Figure S1). (D-O) Confocal z-projections of the indicated genotypes and age, labeled with membrane bound GFP (mCD8-GFP; CD8) driven by the γ specific Gal4 driver GMR71G10-Gal4 (γ-Gal4). While γ-axons of control flies project through the entire lobe (D; n=12/12, F; n=14/14), knockdown of *dpr12* by RNAi (E; n=12/12) or knockout by tsCRISPR (G; n=14/14) resulted in short axons. At L3, γ-axons in *dpr12*^Δ*50-81*^ homozygote mutant animals (L; n=20/20) resemble WT γ-axons (H; n=20/20). At 48h APF, γ-axons normally (I; n=12/12) re-extend to form the adult lobe. *dpr12*^Δ*50-81*^ γ-axons (M; n=14/14) fail to extend to the end of the lobe. This defect persists to adult (J; n=11/11, N; n=18/18). Expressing a *UAS-Dpr12* transgene within γ-KCs in *dpr12*^Δ*50-81*^ homozygote mutant animals rescued the axon regrowth defect (O; n=23/24, K; n=14/14). (P) Quantification of the regrowth defects in J, N and O. The z-projections were blindly classified into four classes of regrowth defect severity, see Figure S1D for examples. Significance was calculated by Kruskal-Wallis test followed by a Mann-Whitney post-hoc test; ***p<0.001. Asterisks demarcate the distal part of the lobe. Green and white indicate mCD8-GFP. Magenta represents FasII staining. Scale bar is 20µm.

The MB is attractive to study wiring of neural circuits not only due to its complex yet stereotypic nature, but also due to its multi-step development. The larval MB is primarily comprised of γ-KCs which form two axonal lobes (a vertical and medial γ-lobes). These are innervated by MBONs and DANs in distinct zones that are different from the adult pattern of zonation (Rohwedder et al., 2016; Saumweber et al., 2018). Subsequently, the γ-lobes undergo extensive remodeling during metamorphosis, including axon pruning followed by developmental regrowth, (Lee et al., 1999) (Figure 1B) to give rise to the adult medial γ-lobe containing the γ1-5 zones (Figure 1A). We have previously demonstrated that regrowth of the adult γ-lobe is genetically controlled by the nuclear receptor Unfulfilled (UNF) functioning as a ligand dependent transcription factor by mechanisms distinct from initial axon outgrowth (Yaniv et al., 2012). Importantly, while we found that UNF promotes axon regrowth partly via the TOR pathway, it is yet unclear through which mechanisms it promotes targeting, circuitry and sub-zone formation. Here, we exploit detailed expression profile analyses to focus on the Immunoglobulin superfamily (IgSF) proteins as potential mediators of zone formation and circuit wiring within the γ4/5 zones.

## Results

### Dpr12 is required for γ neuron regrowth

To identify potential genes and pathways that mediate axon regrowth and circuit formation, we sequenced the RNA content of WT γ-KCs during development (Alyagor et al., 2018) alongside γ-KCs expressing RNAi targeting UNF, a known protein required for regrowth (Figure 1B). Supplementary Table 1 shows this comparison alongside the previously generated (Alyagor et al., 2018) expression profiles of γ neurons expressing a dominant negative form of the Ecdysone Receptor (EcR^DN^), which is required for pruning (Lee et al., 2000; data is freely available in: https://www.weizmann.ac.il/mcb/Schuldiner/resources). The immunoglobulin superfamily (IgSF) appeared as the protein family most significantly affected by UNF RNAi expression (p=3*10^−29^; analyzed in http://www.flymine.org/). Within the IgSF, the Defective Proboscis Response (Dpr) family stood out as 16 of 21 members were significantly expressed in dynamic patterns in developing γ-KCs (Figure 1C, Supplementary Table 2). We found that the transcription of approximately half of the Dprs (7/16; supplemental Table 2) was significantly reduced in UNF-RNAi expressing flies. Interestingly, the interactions between the Dprs, containing two immunoglobulin (Ig) domains, and the Dpr Interacting Proteins (DIPs), containing three Ig domains, are important for proper development and synaptic connectivity of the *Drosophila* visual system and neuromuscular junction (NMJ, Ashley et al., 2019; Venkatasubramanian et al., 2019; Xu et al., 2018). Based on these data, we focused on the Dprs as potential candidates required for regrowth and circuit formation.

We therefore targeted 8 different Dprs in γ-KCs by RNAi (Figure S1A) based on reagent availability (TRiP lines, https://fgr.hms.harvard.edu/fly-in-vivo-rnai). Seven of these RNAi’s did not affect γ neuron development. At this time we cannot conclude whether these Dprs are indeed not required for γ-KCs development or, alternatively, that the lack of phenotype is due to inherent redundancies of Dpr-DIP interactions (Cosmanescu et al., 2018). In contrast, expressing *dpr12* RNAi in γ-KCs induced a dramatic regrowth defect, where axons did not occupy the distal portion of the lobe (Figures 1D-E, S1A). Interestingly, the expression of *Dpr12* is dramatically reduced in neurons expressing UNF-RNAi at the relevant times for regrowth (Figures 1C, S1A, and Supplementary Table 2), suggesting that UNF might positively regulate *dpr12* transcription. These findings suggest that Dpr12 could promote developmental regrowth and circuit formation as a part of an UNF dependent transcriptional program.

To validate the RNAi results, we next perturbed Dpr12 through tissue specific (ts)CRISPR using Gal4 driven Cas9 expression (Meltzer et al., 2019; Port and Bullock, 2016; Port et al., 2020). tsCRISPR of dpr12 in γ-KCs induced a defect closely resembling the RNAi phenotype (Figure 1F-G). Finally, we used CRISPR/Cas9 technology to generate a *dpr12* loss of function mutant (*dpr12*^Δ*50-81*^, Figure S1B-C). At 3^rd^ instar larva (L3), γ-KCs in *dpr12*^Δ*50-81*^ homozygotes exhibited WT morphology (Figure 1H, L) and subsequently pruned normally (data not shown). These data indicate that Dpr12 is not required for initial axon extension or pruning. In contrast, γ-KCs in *dpr12*^Δ*50-81*^ animals failed to extend to the distal part of the lobe at 48h APF, a time when γ-axons would have normally completed their regrowth (Lee et al., 1999; Rabinovich et al., 2016), and in adult (Figure 1I,J, M, N). Importantly, γ-specific expression of a *Dpr12* transgene significantly rescued the regrowth defect within *dpr12*^Δ*50-81*^ homozygotes (Figure 1K, O, P, ranking examples shown in Figure S1D). These data demonstrate that Dpr12 is required for axon regrowth during metamorphosis and that mutant γ-axons stop prematurely and do not extend into the distal end of the lobe.

### Dpr12 is cell autonomously required for γ-axon regrowth into the γ4/5 zones

To determine whether Dpr12 functions in a cell-autonomous manner, we used the Mosaic Analysis with a Repressible Cell Marker (MARCM) technique to express *dpr12-RNAi* within neuroblast (NB) or single cell (SC) clones. We found that both NB and single γ-KC clones expressing *dpr12* RNAi exhibited normal growth at L3 but failed to fully extend axons at 48hr APF and in adult flies (Figure 2A-L). Based on these results, we conclude that *Dpr12* is cell-autonomously required in γ-KCs for their full developmental regrowth.

**Figure 2.**
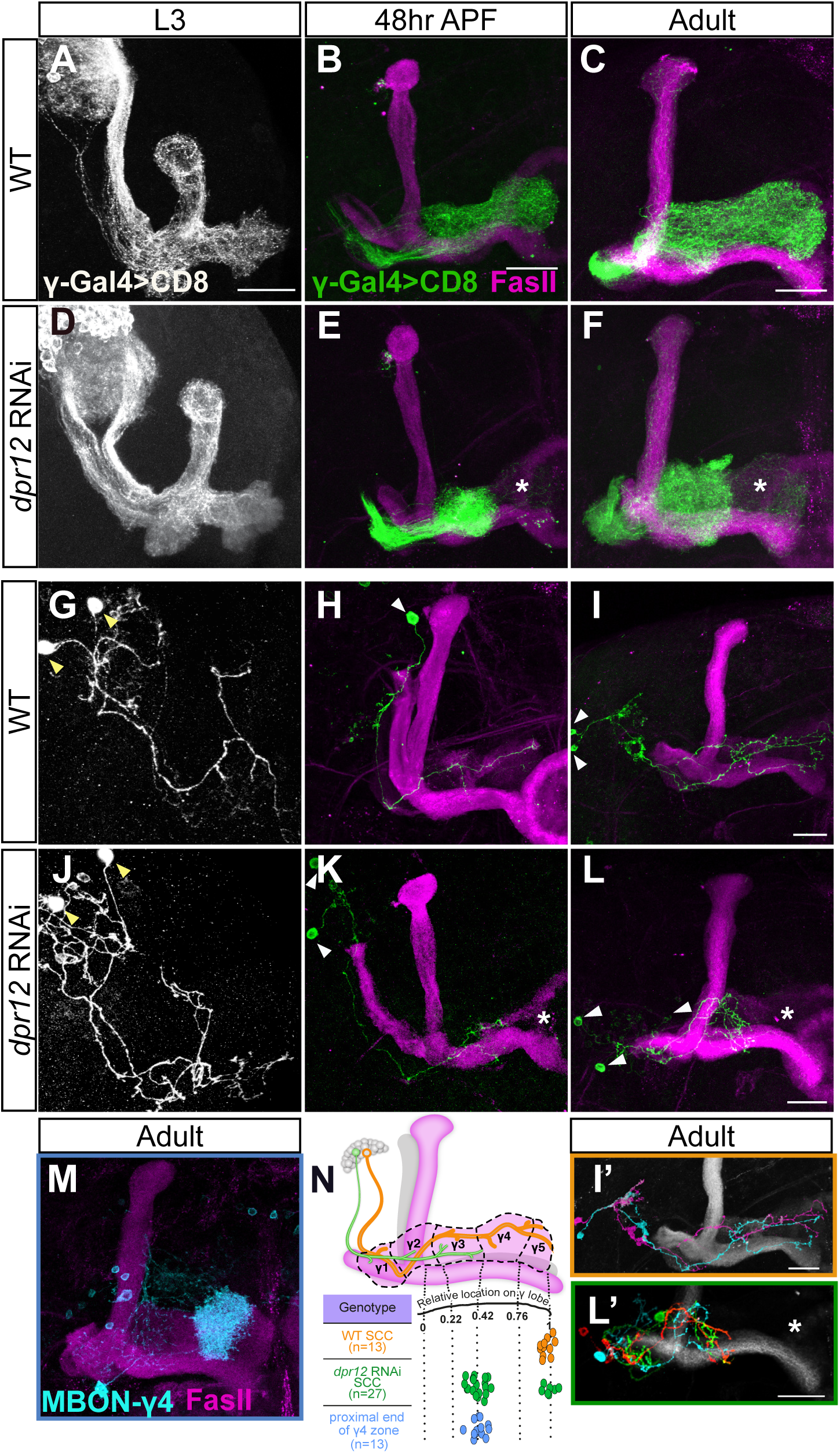
Dpr12 is cell autonomously required for γ-axon regrowth into the γ4/5 zones. (A-L) Confocal z-projections of MARCM neuroblast (NB, A-F) and single-cell (SC, G-L) clones labeled with membrane bound GFP (mCD8-GFP; CD8) driven by the γ specific Gal4 driver GMR71G10-Gal4 (γ-Gal4). At L3, NB and SC clones expressing *dpr12* RNAi are similar to equivalent WT clones (A; n=20/20, D; n=15/15, G; n=15/15 and J; n=17/17). At 48hr APF and adult stage, WT NB (B; n=15/15, C; n=10/10) and SC (H; n=16/16, I; n=13/13) clones extend their axons to form the full adult lobe. In contrast, clones expressing *dpr12* RNAi (E; n=14/14, F; n=22/2, K; n=18/24, L; n=19/27) fail to extend their axons to the distal part of the medial lobe (asterisks). I’ and L’ are traces of the single cell clones depicting each cell in a different color. (M) Confocal z-projection of MBONγ4>γ1γ2 labeled by GMR18H09-Gal4 driving the expression of mCD8-GFP (CD8) shown in cyan. (N) Top: Schematic representation of WT (orange) and *dpr12* RNAi (green) expressing single γ-KC clones. Bottom: Measurements of the relative location to which WT (I) and *dpr12* RNAi (L) axons grow across the entire length of the adult γ lobe alongside the relative position of the proximal end of γ4 zone (M, see also Figure S2). Arrowheads demarcate single cell bodies. Green, white and cyan represent mCD8-GFP. Magenta shows FasII. Scale bar is 20µm.

Interestingly, unlike other mutants that affect developmental regrowth (Yaniv et al., 2012; Yaniv et al., 2020) *dpr12* mutant axons seem to partially regrow but stop prematurely in a particular and stereotypic location along the lobe. Therefore, and given that the γ-lobe is divided into distinct zones, we next mapped the location of the premature stopping in more detail. We measured the length of adult WT and mutant axons relative to the γ-lobe span and superimposed these data onto the γ-lobe zones (Figures 2M, S2), as defined by distinct innervations of MBONs and DANs (Aso et al., 2014a; Shuai et al., 2015). This analysis indicated that the premature stopping of clones expressing *dpr12* RNAi correlates with the border between the γ3 and γ4 zones (Figures 2I’, L’, N, Supplemental movies 1-2). Interestingly, we found that approximately 70% of SC clones stopped at the γ3/4 border, while the remaining clones extended to the end of the lobe (Figure 2N). Since axon stalled at a discrete location, our data suggest that the phenotype does not arise from reduced growth potential, *per se*, but rather through a failure to recognize a molecular signal at a designated and stereotypic location. We conclude that Dpr12 is cell-autonomously required for γ-KC projection into the MB γ4/5 zones.

### Dpr12 and its putative interacting protein DIP-δ localize at the γ4/5 zones

To assess Dpr12 protein localization, we used the *Minos* Mediated Integration Cassette transgene collection to generate a GFP insertion within the endogenous *Dpr12* locus, which should produce a Dpr12-GFP fusion protein (Dpr12GFSTF; Figure S1B-C; Nagarkar-Jaiswal et al., 2015). We found that Dpr12-GFP localized to the distal part of γ-axons at late larva (L3; Figure 3A), then becoming diffuse at 24hr APF, when γ-axons initiate their developmental regrowth (Figure 3B). Finally, at 48h APF and in adulthood, Dpr12 relocalized to the distal part of the lobe (Figure 3C-D) and was restricted to the adult γ4 and γ5 zones (Figure 3D). Our data suggest that Dpr12 is expressed at the right time and place to mediate γ-axon regrowth into the γ4/γ5 zones.

**Figure 3.**
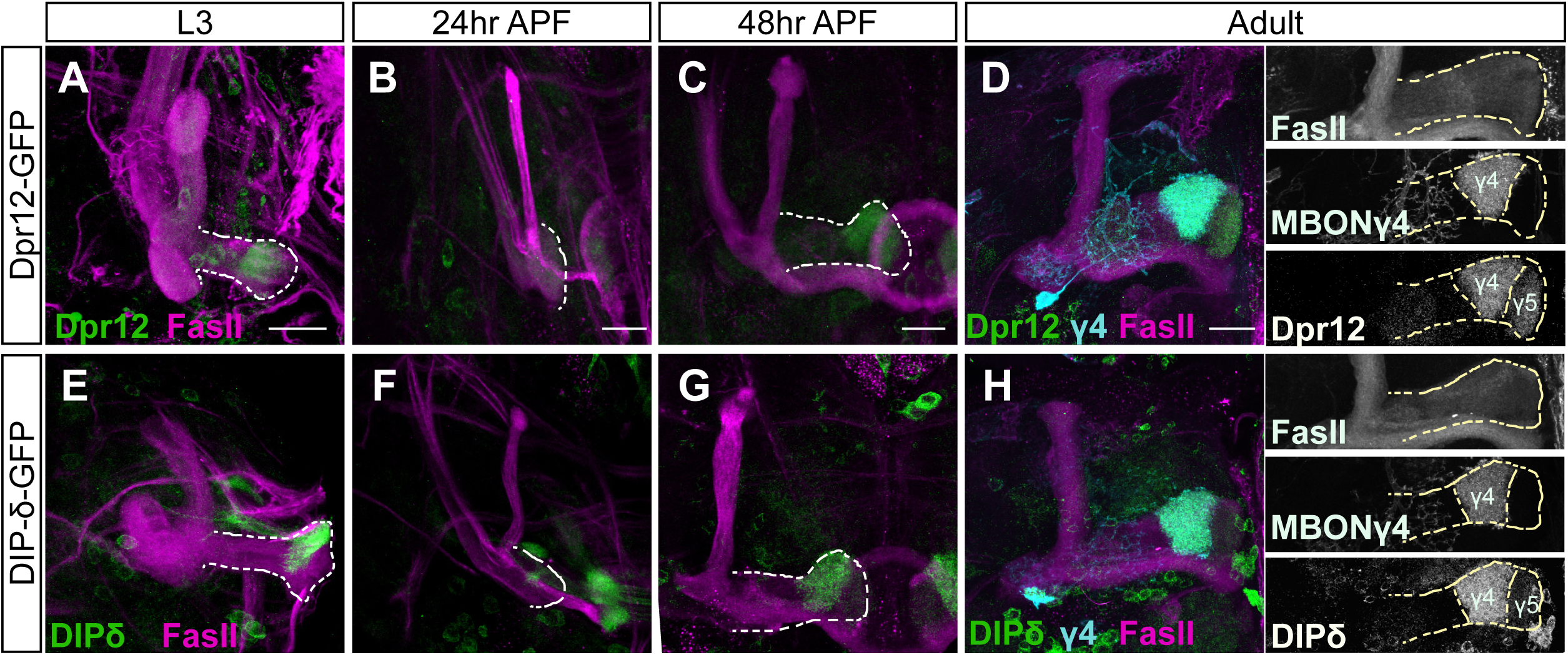
Both Dpr12 and its interacting protein DIP-δ localize to the γ4/5 zones. (A-H) Confocal z-projections of brains expressing MiMIC mediated Dpr12^GFSTF^ (Dpr12**-**GFP) and DIP-δ^GFSTF^ (DIP-δ-GFP) fusion proteins at the indicated time points. See Figures S1 and S3 for more details on the fusion protein structure. (A-D) Dpr12**-**GFP is localized to the distal part of the γ-lobe at L3 (A; n=10/10), 48hr APF (C; n=12/12) and the adult stage (D; n=20/20), where it colocalizes with a γ4 MBON (γ4; labeled by GMR18H09-Gal4 driving the expression of CD4-tdT). At 24hr APF (B; n=10/10), Dpr12**-**GFP appears diffuse. (E-H) DIP-δ-GFP is localized to the distal part of the γ-lobe throughout development: L3 (E; n=16/16), 24hr APF (F; n=10/10), 48hr APF (G; n=12/12) and adult (H; n=24/24). At the adult stage, DIP-δ-GFP is colocalized with a γ4 MBON (γ4). Dashed line depicts the medial γ-lobe. Green is GFP, cyan is CD4-tdT, magenta is FasII. Grayscale in right panels of D and H represent single channels, as marked. Scale bar is 20µm.

Dprs can form heterophilic interactions with DIPs in a rather promiscuous fashion, in which most Dprs can bind to multiple DIPs and most DIPs can bind to multiple Dprs (Carrillo et al., 2015; Cosmanescu et al., 2018; Ozkan et al., 2013). Interestingly, Dpr12 and DIP-δ represent a unique case of ‘monogamous’ binding. We found that DIP-δ-GFP (DIP-δ^GFSTF^; Figure S3A-B) localized, similarly to Dpr12, to the γ4/5 zones (Figure 3H). However, in contrast to Dpr12, DIP-δ was localized to the distal MB lobe at all developmental timepoints tested (Figure 3E-H), including at 24hr APF, when γ-axons are completely pruned and have not yet extended (Figure 3F). Taken together, these data indicate that DIP-δ is localized to γ4/γ5 zones throughout development and is expressed in cells that project to this region before γ-KCs reach their terminal projections.

### DIP-δ is non-cell autonomously required for γ-axon regrowth

We next asked whether DIP-δ is also required for the extension of γ-axons into the γ4/5 zones. We therefore both generated and obtained *DIP-δ* mutant alleles (*DIP-δ*^*T2A-Gal4*^, *DIP-δ*^*1-119*^, respectively; Figure S3A-B). We marked the γ-KCs by expressing a membrane bound Tomato (QUAS-mtdT-3XHA) driven by the Gal4-independent γ-KC specific QF2 driver (71G10-QF2). At L3, γ-KCs within *DIP-δ* mutant brains exhibited WT morphology (Figure 4A, E, S3C), indicating that DIP-δ is not required for their initial axon extension. However, *DIP-δ* mutant brains displayed a γ-axon regrowth defect at 48hr APF, and in adult (Figure 4B, C, F, G, Figure S3D) which resembled the *dpr12* mutant phenotype. To confirm that *DIP-δ* loss-of-function induced this γ-axon regrowth defect, we exploited the fact that the *DIP-δ*^*T2A-Gal4*^ allele also expresses Gal4 in DIP-δ^+^ neurons (Figure S3A-B). Expressing *DIP-δ* transgene driven by *DIP-δ*^*T2A-Gal4*^ rescued the γ-axon extension defect (Figure 4D, H, I, S3E) confirming that DIP-δ is required for γ-axon innervation of the γ4/5 zones.

**Figure 4.**
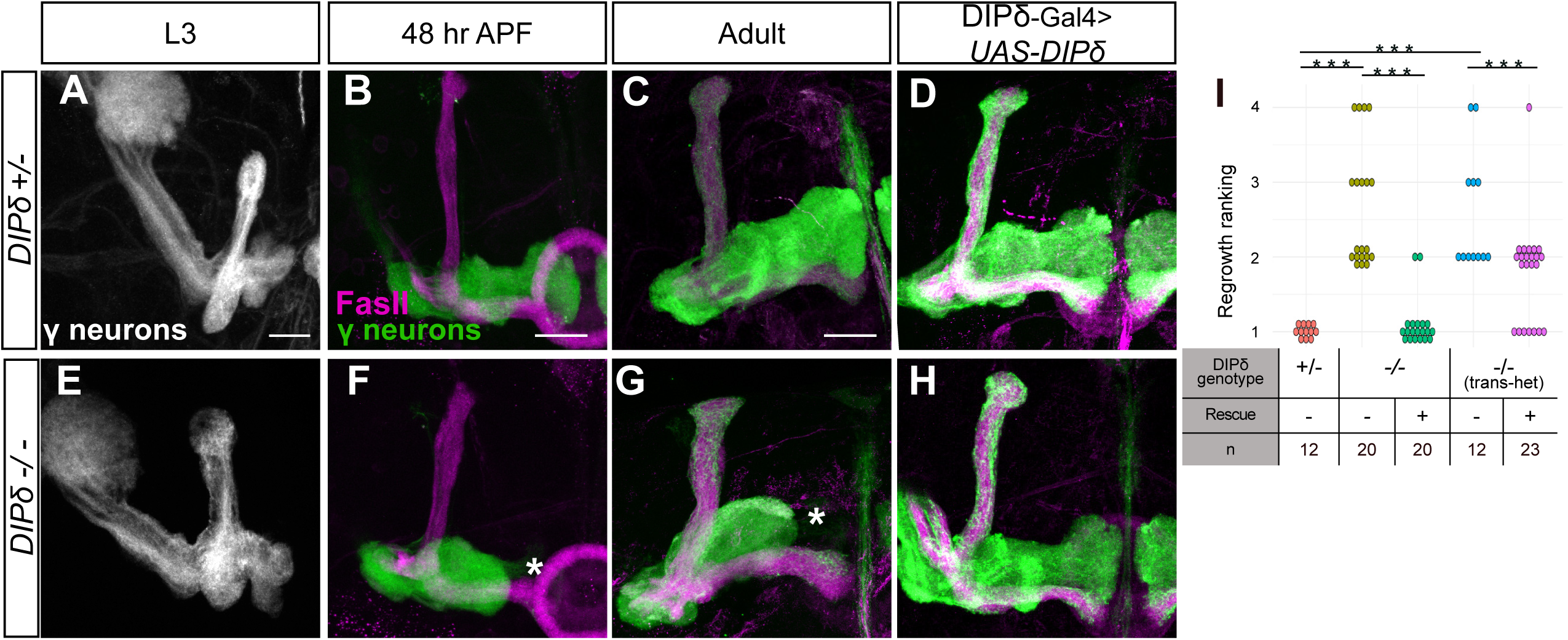
DIP-δ is required for γ-axon regrowth into the γ4/5 zones. (A-H) Confocal z-projections *DIP-δ* hetero- and homozygous brains in which γ-KCs were labeled by expressing membrane bound tandem tomato (mtdT-HA) driven by the γ specific QF2 driver R71G10-QF2 (γ-neurons). Larval (L3) γ-axons grow normally in *DIP-δ*^*T2A-Gal4*^ heterozygotes (A; n=16/16) and homozygotes (E; n=24/24). In contrast, at 48hr APF and adult, γ-axons within *DIP-δ*^*T2A-Gal4*^ homozygotes do not enter the distal part of the lobe (asterisks; F; n=8/8, G; n=20/20), while they grow normally in heterozygotes (B; n=12/12, C; n=12/12). Overexpression of a *DIP-δ* transgene driven by the Gal4 activity of *DIP-δ*^*T2A-Gal4*^ (see also Figure S3) does not affect normal growth (D; n=10/10) and rescues mutant phenotypes (H; n=20/20). Asterisks demarcate distal part of the lobe. Green and white are mtdT-HA, magenta is FasII, Scale bar is 20µm. (I) Quantification of the regrowth defects in C, D, G, H, and Figure S3C-E. Regrowth defect severity and statistics were calculated as in Fig. 1; ***p<0.001

To identify these DIP-δ expressing cells, we expressed DIP-δ RNAi in different cell types, while simultaneously labeling γ-KCs using the QF2 system described above. Driving the expression of *DIP-δ*-RNAi in all glia (using the Pan-glial driver Repo-Gal4) or all KCs (using OK107-Gal4) did not affect γ-axon extension (Figure S4A-B). In contrast, knocking down of *DIP-δ* in all neurons (using the pan-neuronal driver C155-Gal4) or all DIP-δ expressing neurons (using *DIP-δ*^*T2A-Gal4*^) resulted in stalled γ-axons that do not innervate the γ4/5 zones (Figure S4C-D). Similarly, tsCRISPR of *DIP-δ* in all neurons, but not when restricted to γ neurons, affected the extension of γ-axons (Figure S4E-H). Together, these experiments indicate that DIP-δ is not required in γ or other KCs. Rather, DIP-δ likely functions in extrinsic MB neurons in a non-cell autonomous manner to mediate the γ-axon extension into the γ4/5 zones.

### A subpopulation of DANs express DIP-δ in the γ4/γ5 zones

Our data suggest that DIP-δ is expressed in extrinsic MB neurons that innervate the γ4/γ5 zones (Figures 3,4). These zones are strongly innervated by the Protocerebral Anterior Medial (PAM) neurons, a population of about 100 DANs, as well as by specific groups of MBONs (Aso et al., 2014a; Tanaka et al., 2008). To investigate whether DIP-δ is expressed in these cells, we selectively ablated PAM-DANs or MBONs by cell type specific expression of Diphtheria toxin (UAS-DTI) and assayed DIP-δ-GFP localization in adult flies. Ablating one of the γ4 MBONs (MBON-γ4>γ1γ2) did not affect DIP-δ expression (Figure 5A-B). In contrast, ablating PAM DANs using R58E02-Gal4 drastically reduced γ4/5 specific DIP-δ expression (Figure 5C-D). While we cannot exclude the possibility that DIP-δ is additionally expressed in the other γ4 MBON (MBON-γ4γ5), these results strongly suggest that at the adult stage, DIP-δ protein that is localized to the γ4/5 compartments is mainly, if not exclusively, expressed by PAM DANs, consistent with recent profiling experiments (Croset et al., 2018).

**Figure 5.**
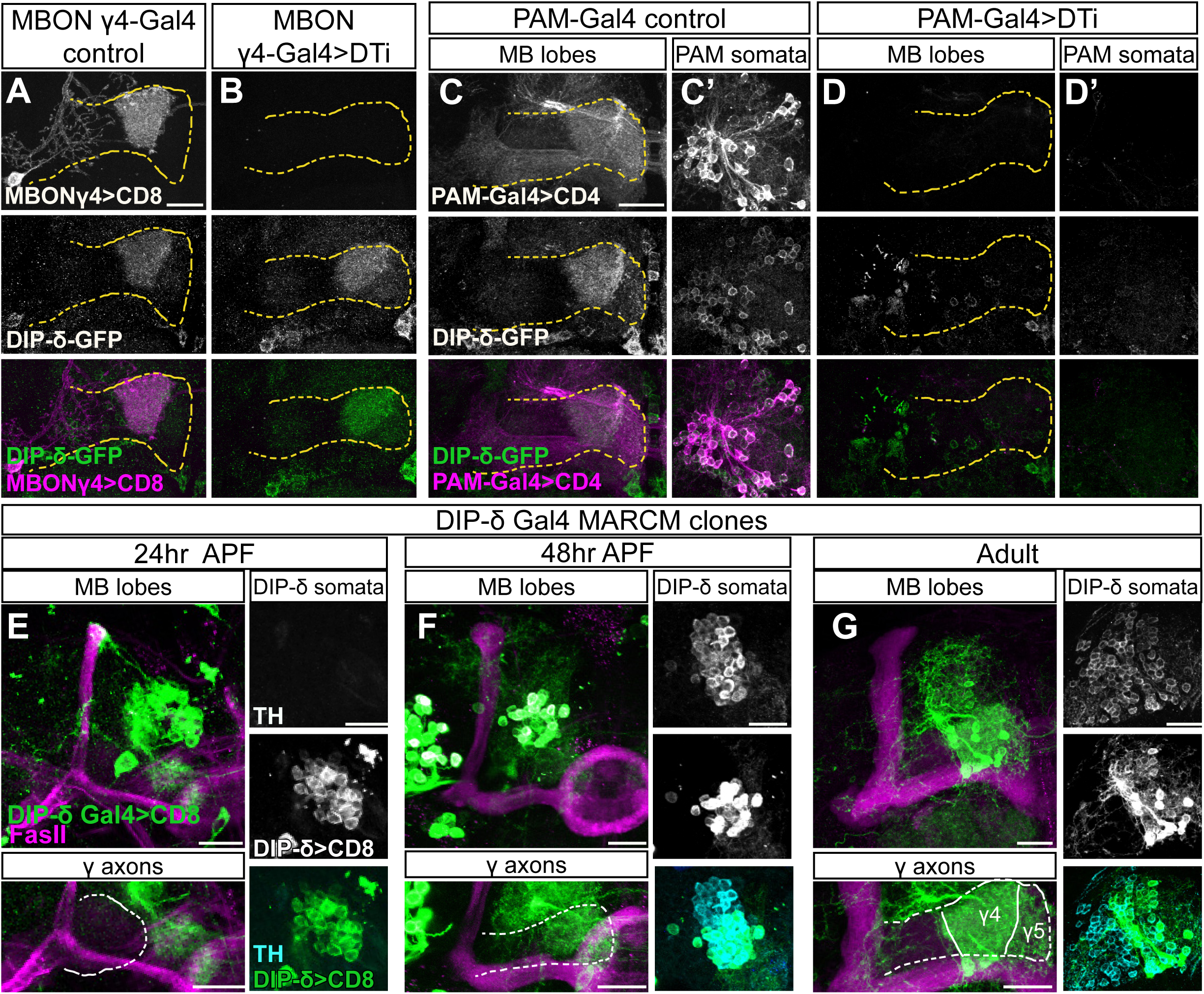
PAM DANs are the source of DIP-δ in the γ4/5 zones. (A-D) Confocal z-projections of brains expressing DIP-δ^GFSTF^ (DIP-δ-GFP) together with the indicated Gal4s and transgenes. Expressing diphtheria toxin (DTi) and the membrane bound RFP (mCD8-RFP; CD8) driven by the γ4 MBON driver MB294B-Gal4 (MBONγ4-Gal4) did not affect DIP-δ-GFP expression (A, n=16/16; B, n=14/14). In contrast, similar expression of DTi and membrane bound Tomato (CD4-tdT; CD4) in PAM-DANs (using the GMR58E02-Gal4; PAM-Gal4) abolished the normal DIP-δ-GFP expression in the γ4/5 zone (compare D, n=18/18, to C, n=16/16) and within the PAM cell bodies (compare D’ to C’). Magenta is CD8-RFP (A-B), CD4-mtdT (C-D). Green is GFP, grayscale depict individual channels as labeled. Scale bar is 20µm. (E-G) Confocal z-projections of MARCM clones labeled by *DIP-δ*^*T2A-Gal4*^ (DIP-δ Gal4) driving the expression of membrane bound GFP (mCD8-GFP; CD8) and heat shocked at 24hr after egg laying. Clones innervate the γ4/5 zones at 24hr APF (E; n=8), 48hr APF (F; n=8) and adult (G; n=16). Clones become tyrosine hydroxylase (TH) positive only at 48hr APF onwards (F,G). Magenta is FasII, green is mCD8-GFP, cyan is TH antibody staining, grayscale single channels are shown as indicated. Scale bar is 20µm.

Next, we visualized PAM-DANs during development to determine if they may provide a template for γ-axon growth, as suggested by DIP-δ-GFP expression (Figure 3). Since the ‘classical’ PAM driver, R58E02, is not expressed throughout development (Figure S5A-C, and data not shown), we analyzed the expression of *DIP-δ*^*T2A-Gal4*^ (Figure S5D-F) and then used the MARCM technique to label sparse *DIP-δ*^*T2A-Gal4*^ clones (Figure 5E-G; of note, these clones remain heterozygous for DIP-δ). We detected DIP-δ expressing clones that innervate the γ4/γ5 zones as early as 24hr APF and up to adulthood. Interestingly, these clones do not seem to express tyrosine hydroxylase (TH), required for Dopamine biogenesis, at 24hr APF but become TH positive at 48hr APF onwards. In summary, our data suggest that DIP-δ is expressed in PAM-DANs, which innervate the future γ4/5 zones as early as 24hr APF, and support the speculation that DIP-δ expressed in PAM-DANs may provide a template for γ-axon growth.

### DIP-δ is required and sufficient for Dpr12 localization

Both Dpr12 and DIP-δ localize to the γ4/5 zones and are required for their formation likely by mediating interactions between the γ-KCs and the PAM-DANs. We therefore investigated how losing either *dpr12* or *DIP-δ* affects their binding partner localization and mature circuit architecture. First, we investigated whether the highly localized expression of Dpr12 and DIP-δ is cell-autonomous or requires interaction with their binding partner. We visualized Dpr12- and DIP-δ-GFP fusion proteins in brains homozygous mutant for their reciprocal Dpr/DIP partner. We found that Dpr12 expression appears diffuse in *DIP-δ* mutant brains throughout development (Figure 6A-D), indicating that Dpr12 protein localization requires interaction with DIP-δ, likely on PAM-DANs. In contrast, DIP-δ localization did not dramatically change during the early stages of development in *dpr12* mutants (Figure 6E-F). At 48hr APF, while we still detected DIP-δ at the distal part of the lobe, it occupied a smaller area than in WT brains and resembled the innervation pattern typical of 24hr APF (compare Figure 6G to Figure 3G). Furthermore, we did not detect any DIP-δ in the adult MB medial lobe (Figure 6H), suggesting that Dpr12 is required for the refinement and maintenance of DIP-δ localization.

**Figure 6.**
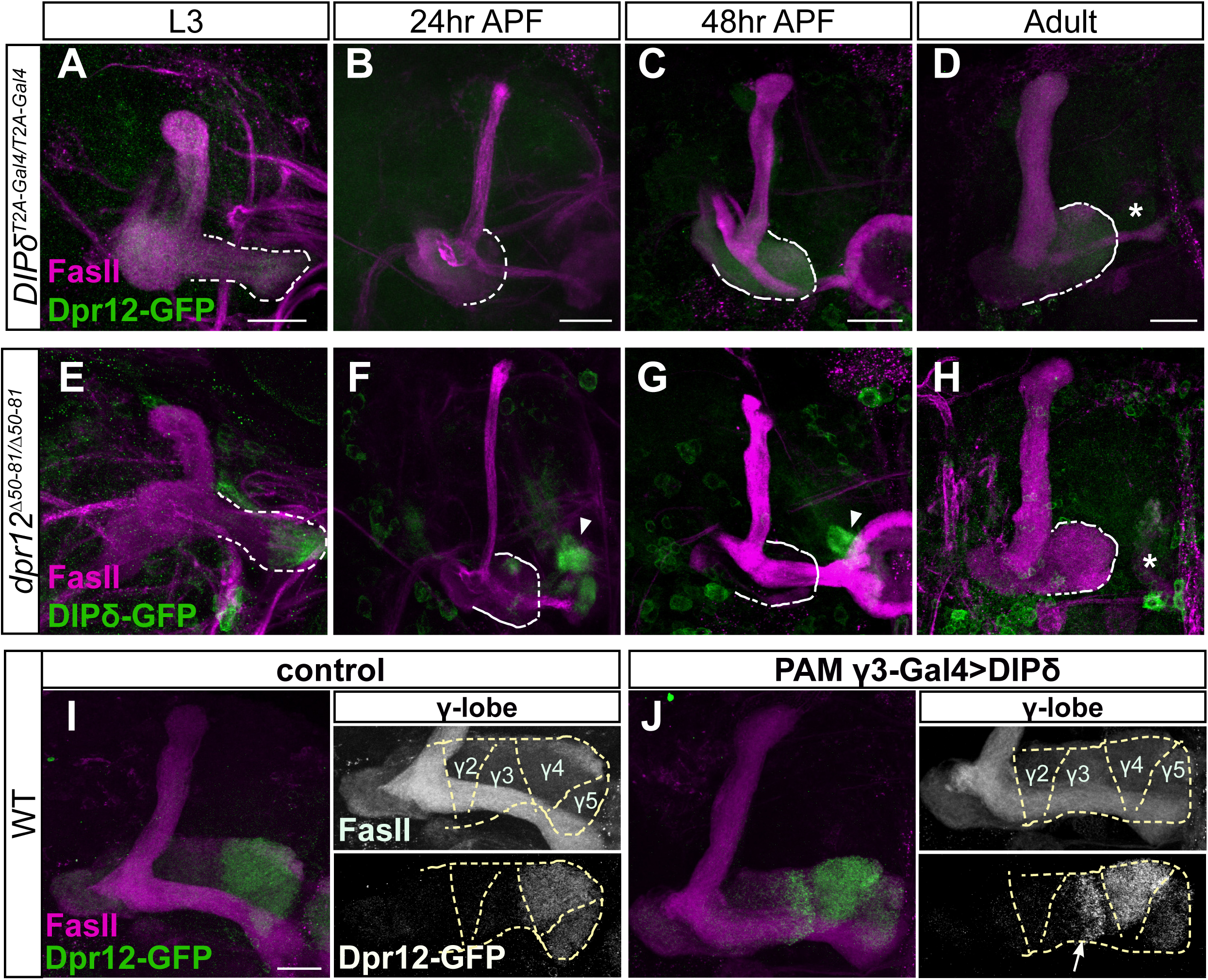
DIP-δ is required and sufficient for Dpr12 localization. (A-J) Confocal z-projections of brains expressing MiMIC mediated Dpr12^GFSTF^ (Dpr12**-**GFP) and DIP-δ^GFSTF^ (DIP-δ-GFP) fusion proteins at the indicated genotypes time points. (A-D) Dpr12**-**GFP expression is diffuse in *DIP-δ*^*T2A-Gal4*^ homozygotes mutant brains at L3 (A; n=20/20), 24hr APF (B; n=14/14), 48hr APF (C; n=28/28) and adult (D; n=26/26). (E-H) DIP-δ-GFP expression in *dpr12*^Δ*50-81*^ homozygotes mutant brains remains localized to the distal part of the γ-lobe at L3 (E; n=16/16), 24hr APF (F; n=16/16), and 48hr APF (G; n=10/10) but cannot be identified in adult brains (H; n=16/16). (I-J) Dpr12**-**GFP expression in WT animals (I, n=8/8) or in those ectopically expressing *DIP-δ* in PAM-DANs that innervate the γ3 zone (J, n=14/14) driven by MB441B-Gal4 (PAMγ3-Gal4). *DIP-δ* expression in PAMγ3 resulted in Dpr12-GFP localization within the γ3 zone (arrow), in addition to its normal γ4/γ5 localization. Arrowheads demarcate DIP-δ expression at the distal part of the lobe. Asterisks demarcate the distal part of the lobe. Dashed line depicts the medial γ-lobe, as determined by FasII staining. Green is GFP, magenta is FasII. Grayscale in right panels of I and J represent single channels, as marked. Scale bar is 20µm.

To explore whether DIP-δ expression is sufficient to regulate Dpr12 protein localization, we mis-expressed DIP-δ in DANs that innervate γ3 zone, normally devoid of Dpr12 protein (Figure 6I). Remarkably, we found that DIP-δ misexpression indeed caused Dpr12 protein to become localized to the adult γ3 zone, in addition to its endogenous γ4/γ5 expression (Figure 6J). Together, these results indicate that while DIP-δ is both required and sufficient for Dpr12 localization throughout development, Dpr12 is required only for maintenance of the adult localization of DIP-δ.

### Dpr12-DIP-δ interaction mediates circuit re-assembly

We next determined if the loss of normal Dpr12-DIP-δ interaction induced axonal misrouting, cell loss or other circuit reorganizations. In *dpr12* mutants, we found that PAM-DANs which normally target the γ4 or γ5 zones misrouted and failed to form substantial connections within the γ-lobe (Figure 7A-D, Figure S6). Importantly, the number of PAM-DAN cell bodies of the subtypes tested remained unchanged (Figure S6C, F). In contrast, γ4-MBON still innervated the γ-lobe in *dpr12* mutant brains, albeit in abnormal locations like the γ3 compartment (Figure 7E-F, Figure S6). Finally, we examined the global neuropil structure in *dpr12* mutant brains by following staining of the active zone protein bruchpilot (Brp), which demonstrated that the γ4/5 zones were largely missing (Figure 7G-H, Movies S3-4). Interestingly, the lack of these domains was accompanied by enlarged γ2/3 zones, as well as distortions in other brain regions (see Crepine, for example, in Figure 7G-H).

**Figure 7.**
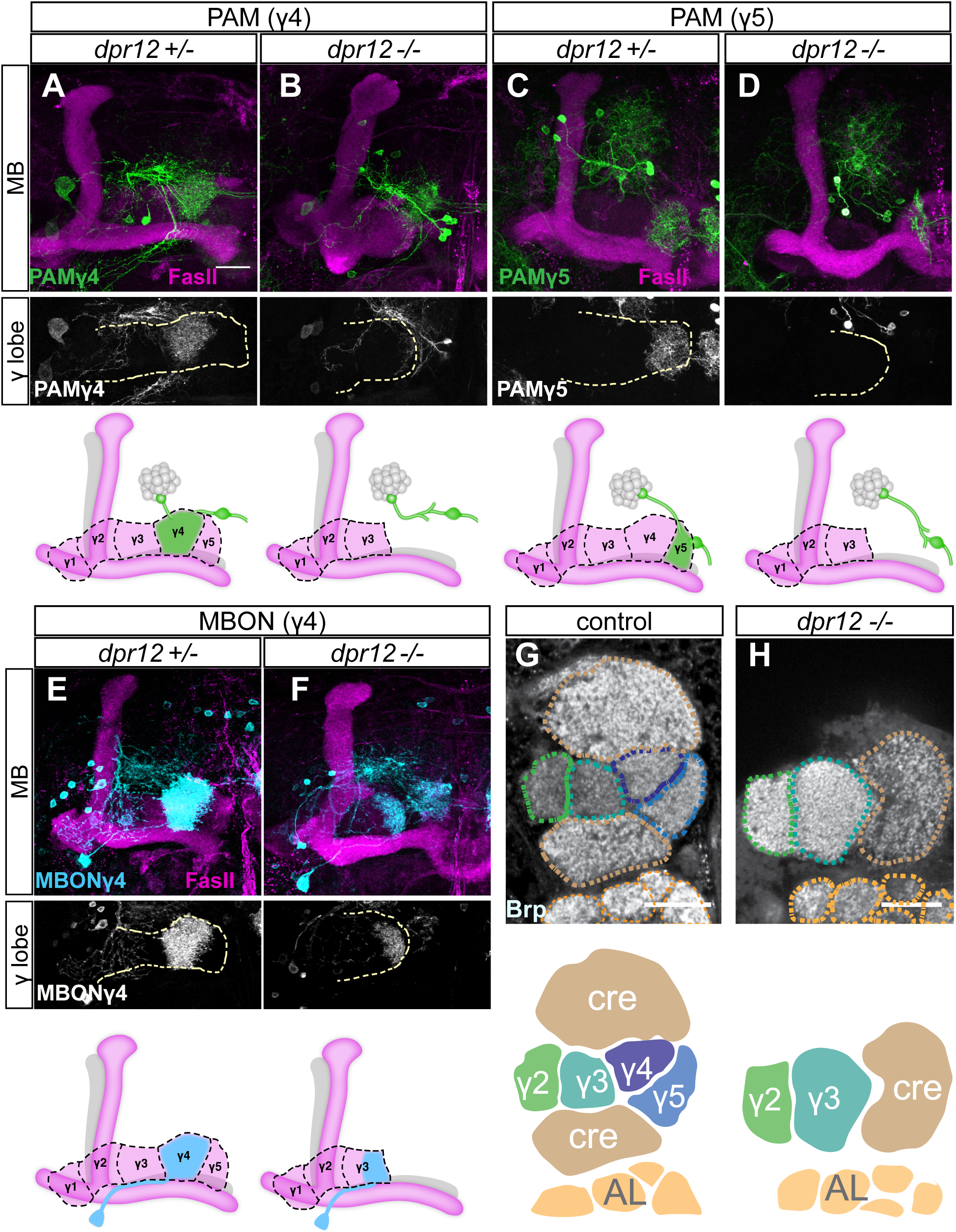
Dpr12-DIP-δ interaction mediates circuit assembly. (A-F) Top: Confocal z-projections of *dpr12*^Δ*50-81*^ heterozygous (A, n=15; C, n=10; E, n=10) and homozygous brains (B, n=8; D, n=18; F, n=24) expressing mCD8-GFP (CD8) driven by: (A-B) R10G03-Gal4 (PAMγ4-Gal4); (C-D) R48H11-Gal4 (PAMγ5-Gal4), or (E-F) R18H09-Gal4 (MBONγ4-Gal4). Bottom: Cartoons schematizing MB lobe structure and innervation by specific PAM-DANs or MBON. (G-H) Single confocal slices of WT (G, n=5/5) and *dpr12*^Δ*50-81*^ homozygous brains (H, n=5/5) stained with anti-Brp. Dashed lines demarcate neuropil boundaries, schematic shown below. cre, crepin; AL, antenna lobe. Magenta is FasII, green and cyan are mCD8-GFP, grayscale depicts single channels as indicated. Scale bar is 20µm.

Taken together, our results suggest that the Dpr12-DIP-δ interaction is required for γ4/5 zone formation by mediating interactions between γ-KCs and a subpopulation of PAM-DANs, eventually instructing their adult-specific connectivity.

## Discussion

Our understanding of the development of complex neural circuits remains largely unknown. Specifically, how long axons can make *en passant* synapses with different partners in a stereotypic manner is not well understood. The unique development and morphology of the *Drosophila* MB γ-lobe, combined with the awesome genetic power of the fly, offers an excellent opportunity to dissect mechanisms required for wiring of complex neural network and specifically for mechanisms that drive zonation within axonal bundles to allow for stereotypic localized innervation by distinct populations of neurons. Here, we identify a molecular mechanism that mediates neuron-neuron interactions that subsequently promote the formation of stereotypic circuits that define subcellular axonal zones.

The adult γ-lobe is divided into zones (also known as compartments) due to specific and localized innervations by extrinsic MB neurons including MBON and DANs. Here we show that the interaction between two IgSF proteins, Dpr12 on γ-KCs and DIP-δ on PAM-DANs, underlies the formation of MB γ4/5 zones. Within each zone, input from DANs can modify synaptic strength between the KC and MBON to provide specific valence to sensory information(Aso et al., 2014a; Aso et al., 2014b; Cognigni et al., 2018; Cohn et al., 2015). Based on the results presented here, we speculate that various specific combinations of adhesion molecules may mediate target recognition events that occur between predefined pre- and post-synaptic pairs in other zones as well. γ-neurons express a broad spectrum of IgSFs in tight temporal regulation (Supplementary Table 1; Alyagor et al., 2018), highlighting their potential role in circuit formation. However, many adhesion molecules, including Dpr/DIPs, can form promiscuous interactions, making their analyses challenging. Future studies could use CRISPR/Cas9 technology to generate multi-gene mutations to further explore the adhesion code required for zone/compartment formation.

Here we used the interaction between Dpr12 and DIP-δ to study the development of the γ4/5 zones. Our developmental analyses have concluded that DIP-δ expressing PAM-DANs arrive to the region of the γ4/5 zones long before γ-axons. Interestingly, in *dpr12* mutant animals PAM-DANs arrive to the right place, linger there for a while (∼48h APF) and then eliminate their γ4/5 innervations, while maintaining and even strengthening/broadening other connections in the MB vicinity. Therefore, it is attractive to speculate that γ-axons extend into a prepatterned lobe. More studies comparing the development of other compartment-specific DANs as well as MBONs are however required.

Here we demonstrate that Dpr12 is cell-autonomously required in γ-KCs while DIP-δ is required on PAM-DANs for the formation of the γ4/5 zones. To the best of our knowledge, this is the first case in which a Dpr molecule was shown to be cell-autonomously required for correct wiring. However, many unresolved questions remain: 1) Why do the γ-axons stop prematurely? One possibility is that axon growth into the γ4/5 zones depends on Dpr12-DIP-δ interaction either because they overcome a yet undiscovered inhibitory signal or because they are positively required for the progression of the growth cone. Alternatively, Dpr12-DIP-δ interaction could be important for the stabilization of the connections between γ-axons and DAN processes to result in the formation of the γ4/5 zones. At 48h APF, the large majority of *dpr12* mutant γ-axons do not innervate the γ4/5 zones, arguing against the stability hypothesis; 2) What are the signaling pathways that mediate Dpr/DIP targeting recognition? None of the Dprs or DIPs contain a large intracellular domain that is capable of signaling. Identifying the co-receptor/s is a critical step in gaining a mechanistic understanding of axon targeting whether in the visual, motor or MB circuits. The Dpr12/DIP-δ interaction which we have uncovered to be required for the formation of specific MB zones is an attractive model to further delve into these mechanisms due to its robust phenotype; 3) What is the significance of the GPI anchor? Many of the Dprs and DIPs, including DIP-δ (data not shown) are predicted GPI anchored proteins, suggesting that they can be cleaved to create a secreted soluble form. Whether this is an important step in targeting has not yet been investigated. Interestingly, the vertebrate homologs of the DIPs, the IgLON subfamily (Zinn and Ozkan, 2017), are GPI anchored proteins that were shown to be cut by metalloproteinases to promote axonal outgrowth (Sanz et al., 2015).

Expression patterns of Dpr and DIP molecules in the neuromuscular junction (NMJ) (Carrillo et al., 2015) and visual system (Carrillo et al., 2015; Tan et al., 2015) suggested a model where these molecules instruct target cell specificity. Recent loss-of-function experiments strengthened this target specificity hypothesis, as the DIP-α-Dpr10 interaction was shown to be important for motoneuron innervation of specific larval (Ashley et al., 2019) and adult (Venkatasubramanian et al., 2019) muscles, and DIP-α-Dpr10/Dpr6 interaction for specific layer targeting in the visual system (Xu et al., 2018). Our results suggest that mechanisms used to target axons and dendrites to specific cell types or layers may be further implicated to orchestrate the wiring of long axons to different pre- and post-synaptic partners along their route and thus the formation of axonal zones.

Here we describe that interaction between two IgSF proteins mediates transneuronal communication that is required for proper wiring within specific zones of the *Drosophila* MB. The anatomical organization of the MB suggests that these interactions may provide target specificity for the long KC axon while it forms *en passant* synapses with different targets along its length. While the existence of such wiring architecture is known from invertebrates such as *Drosophila* and *C. elegans*, long axons making distinct yet stereotypic *en passant* connections is not widely described in vertebrates. Given the existence of long axons, that travel through dense neuropil structures, such as mossy fibers in the hippocampus, cholinergic axons in the basal forebrain, and parallel fibers in the cerebellar cortex, we posit that this type of connectivity exists in vertebrates but has not yet been described in detail due to technological limitations that are likely to be resolved soon. Pairwise IgSF molecular interactions are conserved in vertebrates and invertebrates, implying similar mechanisms to dictate axon and dendrite targeting of subcellular neurite zones in other organisms.

## Acknowledgments

We thank Larry Zipursky for sharing unpublished reagents, the Bloomington Stock Centers for reagents; monoclonal antibodies were obtained from the Developmental Studies Hybridoma Bank developed under the auspices of the NICHD and maintained by the University of Iowa. We thank R. Rothkopf for assistance with statistics; M. Schuldiner, A. Yaron, T. Misgeld and the O.S lab for discussions and critical reading of this manuscript. We thank Life Science Editors for editing assistance. Funding: This work was supported by the European Research Council (erc), consolidator grant # 615906, “AxonGrowth” and the Volkswagen Stiftung (joint Lower Saxony – Israel) grant # ZN3459. Fly food for this project was funded by the Women Health Research Center. O.S. is the Incumbent of the Prof. Erwin Netter Professorial Chair of Cell Biology, T.R. was supported by Axa as team member of the Axa Chair from genome to structure, F.R is a GSfBS member and was supported by Evangelisches Studienwerk Villigst e.V. Competing interests: Authors declare no competing interests. Data and materials availability: All data is available in the main text or the supplementary materials. In addition, RNAseq data will be available via Gene Expression Omnibus at NCBI.

## Supplemental materials and STAR methods

**KEY RESOURCES TABLE**

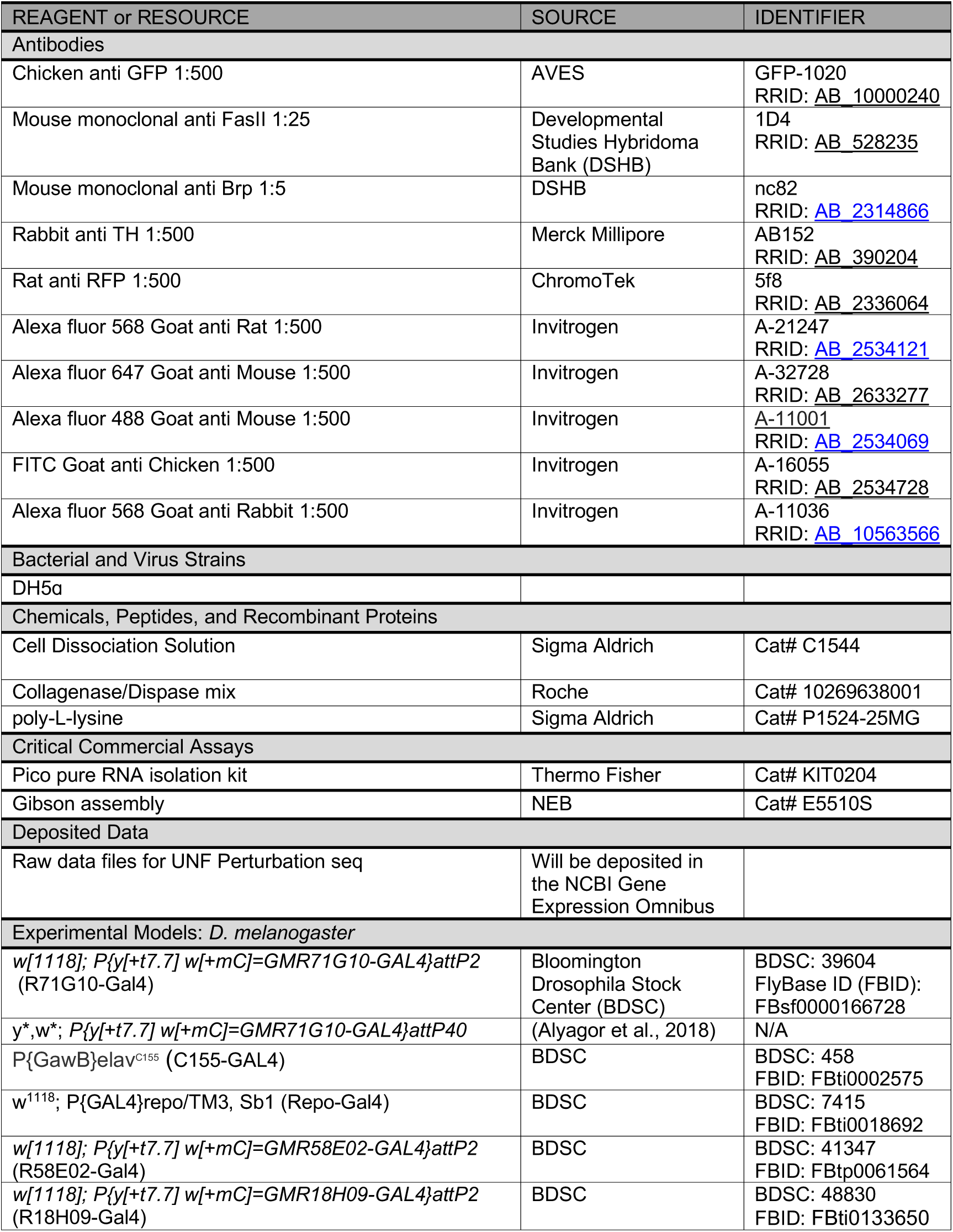
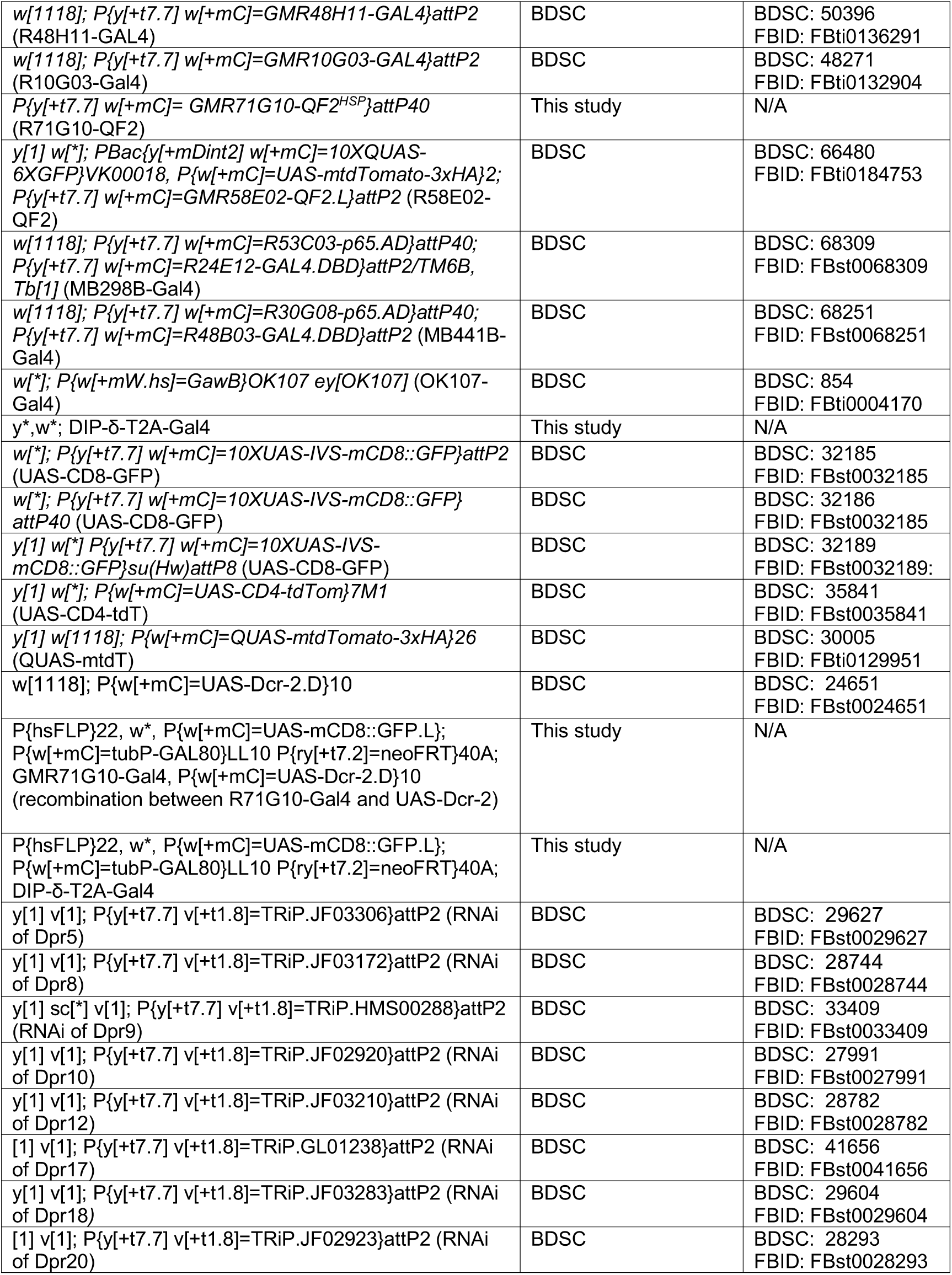
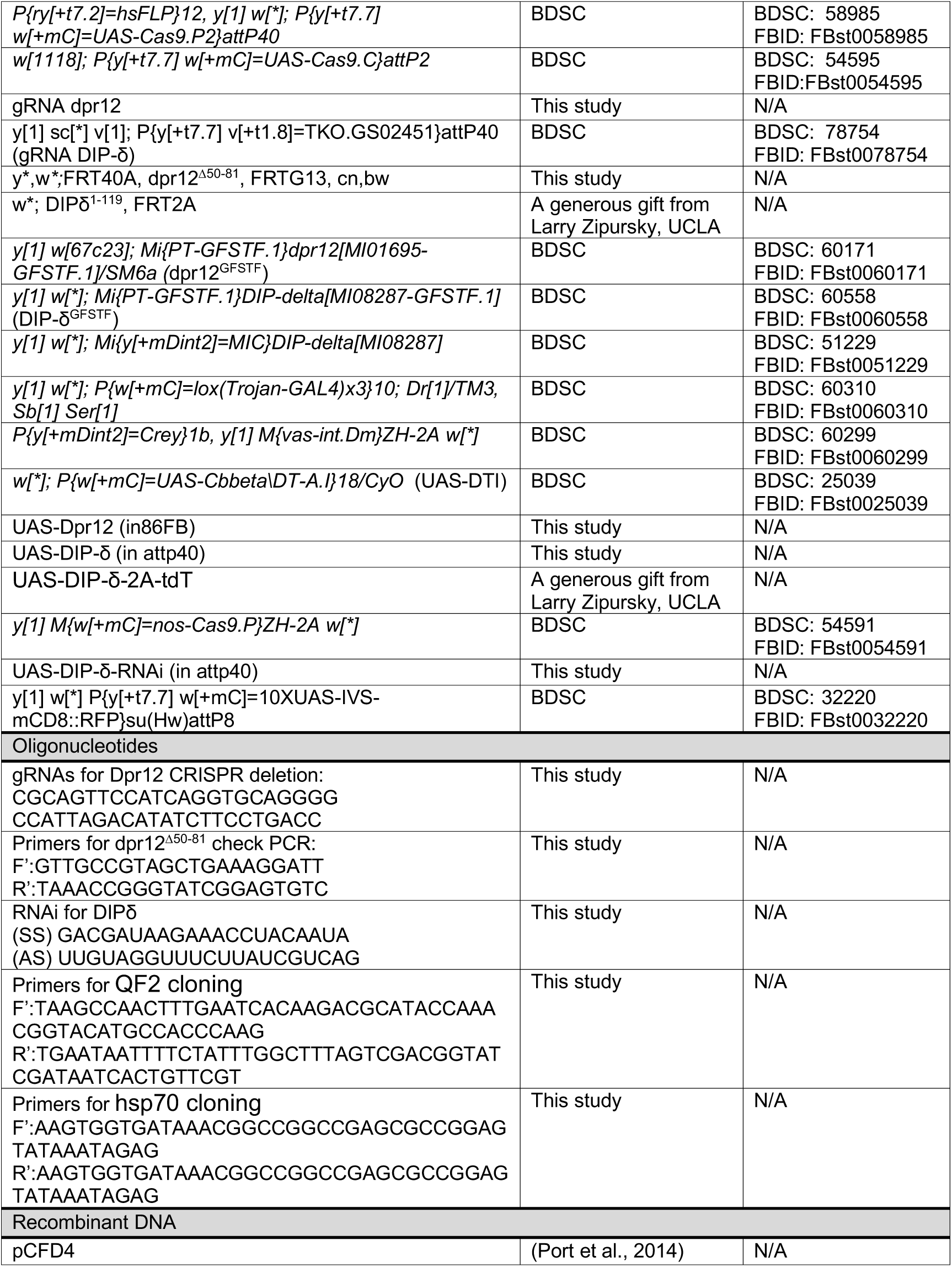
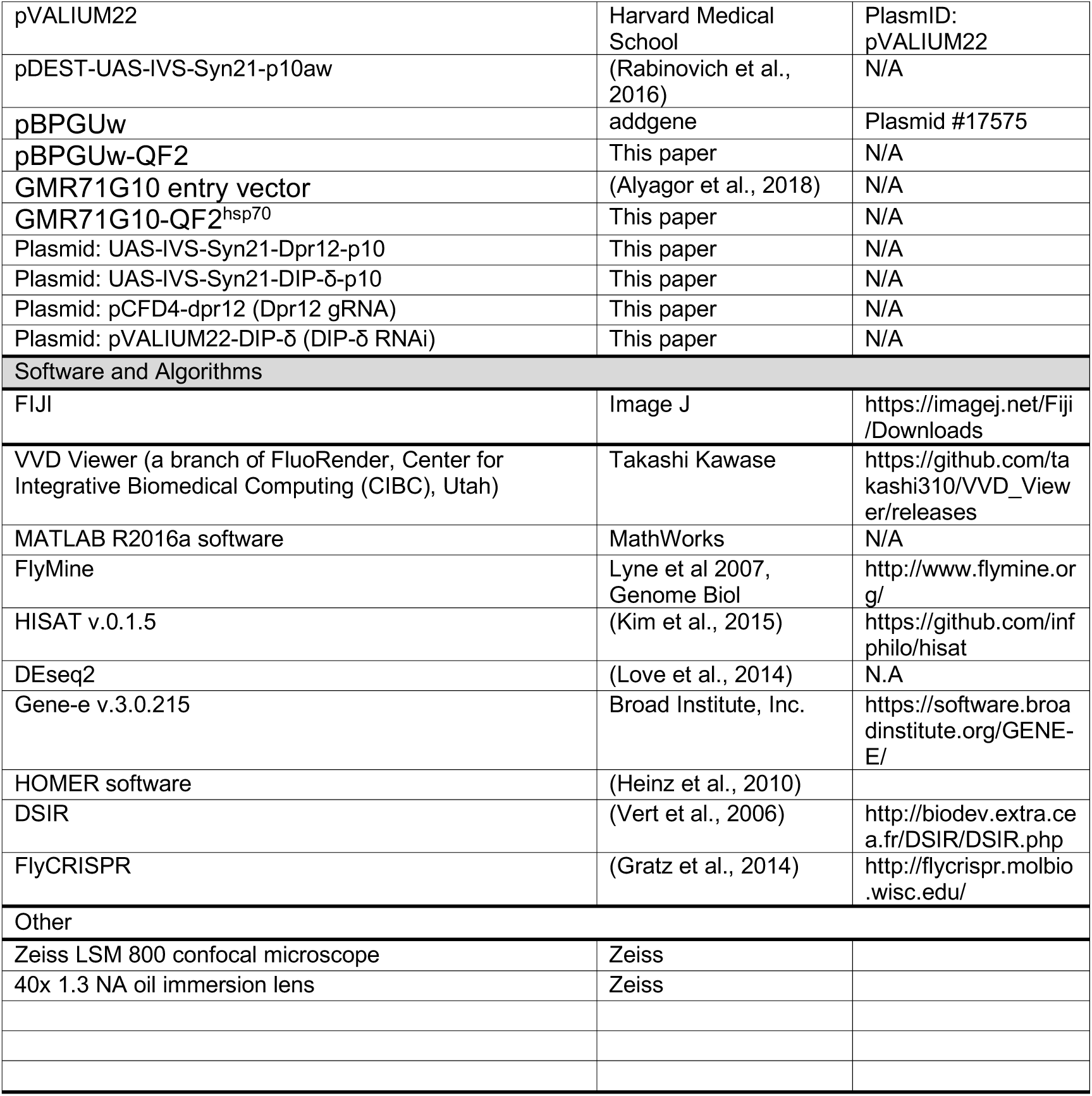

## CONTACT FOR REAGENT AND RESOURCE SHARING

Further information and requests for resources and reagents should be directed to and will be fulfilled by the Lead Contact, Oren Schuldiner

## EXPERIMENTAL MODEL

*Drosophila melanogaster* flies were reared under standard laboratory conditions at 25°C on molasses containing food. Males and females were chosen at random. For developmental analysis white pupae were collected and incubated for the indicated number of hours. For adult analysis flies were dissected 3-5 days post eclosion. For a detailed list of the stocks and their source, see Key Resource Table.

## GENOTYPES

hsFLP is y,w,hsFLP122; GFP is GFP; tdT is tdTom; mtdT-HA is mtdTomato-3xHA; 40A and G13 are FRTs on 2L and 2R respectively; R71G10 is GMR71G10-Gal4; R18H09 is GMR18H09-Gal4; R58E02 is GMR58E02-Gal4; R10G03 is GMR10G03-Gal4; R48H011 is GMR48H011-Gal4; G80 is TubP-Gal80. Males and females were used interchangeably but only the female genotype is mentioned.

**Figure 1**

(D) y, w; R71G10, UAS-mCD8-GFP/+; R71G10, UAS-Dcr-2/+

(E) y, w; R71G10, UAS-mCD8-GFP/+; R71G10, UAS-Dcr-2/ TRiP.JF03210(dpr12-RNAi)

(F) y, w; R71G10, UAS-mCD8-GFP/+; UAS-Cas9.C

(G) y, w; R71G10, UAS-mCD8-GFP/+; UAS-Cas9.C/dpr12-gRNA

(H-J) y, w; UAS-mCD8-GFP/+; R71G10/+

(K) y, w; UAS-mCD8-GFP/+; R71G10/UAS-Dpr12

(L-N) y, w; 40A, dpr12^Δ50-81^, G13/40A, dpr12^Δ50-81^, G13; R71G10, UAS-mCD8-GFP/+

(O) y, w; 40A, dpr12^Δ50-81^, G13/40A, dpr12^Δ50-81^, G13; R71G10, UAS-mCD8-GFP/UAS-Dpr12

**Figure 2**

(A-C, G-I) hsFLP, UAS-mCD8-GFP; Gal80, FRT40A/FRT40A; R71G10, UAS-Dcr-2/+ (D-F, J-L) hsFLP, UAS-mCD8-GFP; Gal80, FRT40A/FRT40A; R71G10, UAS-Dcr-2/ TRiP.JF03210 (dpr12-RNAi)

(M) y, w; UAS-mCD8-GFP/+; R18H09/+

**Figure 3**

(A-C) y, w; Dpr12 ^GFSTF^/+

(D) y, w; UAS-CD4-tdT/Dpr12 ^GFSTF^; R18H09/+

(E-G) y, w; DIP-δ ^GFSTF^/+

(H) y, w; UAS-CD4-tdT/+; DIP-δ ^GFSTF^/R18H09

**Figure 4**

(A-C) y, w; R71G10-QF2, QUAS-mtdT-HA/+; *DIP-δ*^*T2A-Gal4*^/+

(D) y, w; R71G10-QF2, QUAS-mtdT-HA/UAS-*DIP-δ*; *DIP-δ*^*T2A-Gal4*^/+

(E-G) y, w; R71G10-QF2, QUAS-mtdT-HA/+; *DIP-δ*^*T2A-Gal4*^/*DIP-δ*^*T2A-Gal4*^

(H) y, w; R71G10-QF2, QUAS-mtdT-HA/UAS-*DIP-δ*; *DIP-δ*^*T2A-Gal4*^/*DIP-δ*^*T2A-Gal4*^

**Figure 5**

UAS-mCD8-RFP; R53C03-p65.AD/+; DIP-δ ^GFSTF^/R24E12-GAL4.DBD

UAS-mCD8-RFP; R53C03-p65.AD/UAS-DTi; DIP-δ ^GFSTF^/R24E12-GAL4.DBD

y, w; UAS-CD4-tdT/+; DIP-δ ^GFSTF^/R58E02

y, w; UAS-CD4-tdT/UAS-DTi; DIP-δ ^GFSTF^/R58E02

(E-G) hsFLP, UAS-mCD8-GFP; Gal80, FRT40A/FRT40A; *DIP-δ*^*T2A-Gal4*^/+

**Figure 6**

(A-D) y, w; Dpr12 ^GFSTF^/+; *DIP-δ*^*T2A-Gal4*^/*DIP-δ*^*T2A-Gal4*^

(E-G) y, w; 40A, dpr12^Δ50-81^, G13/40A, dpr12^Δ50-81^, G13; DIP-δ ^GFSTF^/+

(I) y, w; Dpr12 ^GFSTF^/+; UAS-DIP-δ-2A-tdT/+

(J) y, w; Dpr12 ^GFSTF^ /R30G08-p65.AD; R48B03-GAL4.DBD/ UAS-DIP-δ-2A-tdT

**Figure 7**

(A) y, w; 40A, dpr12^Δ50-81^, G13/+; R10G03/UAS-mCD8-GFP

(B) y, w; 40A, dpr12^Δ50-81^, G13/40A, dpr12^Δ50-81^, G13; R10G03/UAS-mCD8-GFP

(C) y, w; 40A, dpr12^Δ50-81^, G13/+; R48H011/UAS-mCD8-GFP

(D) y, w; 40A, dpr12^Δ50-81^, G13/40A, dpr12^Δ50-81^, G13; R48H011/UAS-mCD8-GFP

(E) y, w; 40A, dpr12^Δ50-81^, G13/+; R18H09/UAS-mCD8-GFP

(F) y, w; 40A, dpr12^Δ50-81^, G13/40A, dpr12^Δ50-81^, G13; R18H09/UAS-mCD8-GFP

(G) w1118

(H) w^1118^; 40A, dpr12^Δ50-81^, G13/40A, dpr12^Δ50-81^, G13

**Figure S1**

(A)

y, w; R71G10, UAS-mCD8-GFP/+; R71G10, UAS-Dcr-2/ TRiP.JF03306(Dpr5-RNAi)

y, w; UAS-mCD8-GFP/+; R71G10/ TRiP.JF03172 (Dpr8-RNAi)

y, w; UAS-mCD8-GFP/+; R71G10/ TRiP.HMS00288 (Dpr9-RNAi)

y, w; R71G10, UAS-mCD8-GFP/+; R71G10, UAS-Dcr-2/ TRiP.JF02920 (Dpr10-RNAi)

y, w; R71G10, UAS-mCD8-GFP/+; R71G10, UAS-Dcr-2/ TRiP.JF03210(Dpr12-RNAi)

y, w; UAS-mCD8-GFP/+; R71G10/ TRiP.GL01238 (Dpr17-RNAi)

y, w; R71G10, UAS-mCD8-GFP/+; R71G10, UAS-Dcr-2/ TRiP.JF03283}attP2 (Dpr18-RNAi)

y, w; UAS-mCD8-GFP/+; R71G10/ TRiP.JF02923 (Dpr20-RNAi) (D)

y, w; 40A, Dpr12^Δ50-81^, G13/40A, Dpr12^Δ50-81^, G13; R71G10, UAS-mCD8-GFP/+

y, w; 40A, Dpr12^Δ50-81^, G13/40A, Dpr12^Δ50-81^, G13; R71G10, UAS-mCD8-GFP/+

y, w; 40A, Dpr12^Δ50-81^, G13/40A, Dpr12^Δ50-81^, G13; R71G10, UAS-mCD8-GFP/+

y, w; UAS-mCD8-GFP/+; R71G10/+

**Figure S2**

(A) hsFLP, UAS-mCD8-GFP; Gal80, FRT40A/FRT40A; R71G10, UAS-Dcr-2/+

(B) hsFLP, UAS-mCD8-GFP; Gal80, FRT40A/FRT40A; R71G10, UAS-Dcr-2/ TRiP.JF03210 (Dpr12-RNAi)

(D) y, w; UAS-mCD8-GFP/R30G08-p65.AD; R48B03-GAL4.DBD/ +

(E) y, w; UAS-mCD8-GFP/+; R18H09/ +

(F) y, w; UAS-mCD8-GFP/+; R48H011/+

**Figure S3**

(C-D) y, w; R71G10-QF2, QUAS-mtdT-HA/+; *DIP-δ*^*T2A-Gal4*^/ *DIP-δ*^*1-119*^

(E) y, w; R71G10-QF2, QUAS-mtdT-HA/UAS-*DIP-δ*; *DIP-δ*^*T2A-Gal4*^/*DIP-δ*^*T2A-Gal4*^

**Figure S4**

(A) y, w; UAS-*DIP-δ-RNAi*/R71G10-QF2, QUAS-mtdT-HA; Repo-Gal4, UAS-mCD8-GFP/+

(B) y, w; UAS-*DIP-δ-RNAi*/R71G10-QF2, QUAS-mtdT-HA; OK107-Gal4

(C) C155-Gal4; UAS-*DIP-δ-RNAi*/R71G10-QF2, QUAS-mtdT-HA

(D) y, w; UAS-*DIP-δ-RNAi*/R71G10-QF2, QUAS-mtdT-HA; *DIP-δ*^*T2A-Gal4*^/+

(E) C155-Gal4; UAS-Cas9.P2/+

(F) C155-Gal4; UAS-Cas9.P2/TKO.GS02451(*gRNA DIP-δ)*

(G) y, w; R71G10, UAS-mCD8-GFP/ +; UAS-Cas9.C/+

(H) y, w; R71G10, UAS-mCD8-GFP/ TKO.GS02451(*gRNA DIP-δ)*; UAS-Cas9.C/+

**Figure S5**

(A-C) y, w; QUAS-GFP, UAS-mtdT-HA/UAS-tdT; R58E02/R58E02-QF2

(D-F) y, w; R71G10-QF2, QUAS-mtdT-HA/+; *DIP-δ*^*T2A-Gal4*^/+

**Figure S6**

(A) y, w; 40A, dpr12^Δ50-81^, G13/+; R10G03/UAS-mCD8-GFP

(B) y, w; 40A, dpr12^Δ50-81^, G13/40A, dpr12^Δ50-81^, G13; R10G03/UAS-mCD8-GFP

(D) y, w; 40A, dpr12^Δ50-81^, G13/+; R48H011/UAS-mCD8-GFP

(E) y, w; 40A, dpr12^Δ50-81^, G13/40A, dpr12^Δ50-81^, G13; R48H011/UAS-mCD8-GFP

(G) y, w; 40A, dpr12^Δ50-81^, G13/+; R18H09/UAS-mCD8-GFP

(H) y, w; 40A, dpr12^Δ50-81^, G13/40A, dpr12^Δ50-81^, G13; R18H09/UAS-mCD8-GFP

## METHOD DETAILS

### RNA extraction

The RNA extraction of WT and UNF-RNAi expressing MB γ neurons was performed as described in Ref #15. In brief, brains were dissected in a cold Ringer’s solution and dissociated by incubation with collagenase/dispase mix at 29°C (Roche, 15 minutes for larval and pupal brains and 30 minutes for adult brains), washed in dissociation solution (Sigma-Aldrich), and mechanically dissociated into single cells. Cells were transferred via 35 µm mesh (Falcon) to eliminate clusters and debris. 1000 γ neurons (DsRed^+^) were sorted using a 100 mm nozzle and low pressure in BD FACSAria Fusion (BD Bioscience) directly into 100 µl Pico-Pure RNA isolation kit extraction buffer (Life Technologies) followed by RNA extraction. mRNA was captured using 12 ml of Dynabeads oligo (Life Technologies), which were washed from unbound total RNA according to the protocol. mRNA was eluted from beads at 85°C with 10 ml of 10 mM Tris-Cl (pH 7.5). mRNA was barcoded, converted into cDNA and linearly amplified by T7 *in vitro* transcription. The resulting RNA was fragmented and converted into an Illumina sequencing-ready library through ligation, RT, and PCR. Prior to sequencing, libraries were evaluated by Qubit fluorometer and TapeStation (Agilent).

### Analysis of RNA-Seq Data

Samples were sequenced using Illumina NextSeq 500, at a sequencing depth of an average of 5 million reads. We aligned the reads to *D. melanogaster* reference genome (DM6, UCSC) using Hisat v0.1.5 with ‘‘–sensitive -local’’ parameters (Kim et al., 2015). Gene annotation were taken from FlyBase.org (Dmel R6.01/Fb_2014_04). Duplicate reads were filtered if they aligned to the same base and had identical unique molecular identifiers (UMI). Expression levels were counted using HOMER software (http://homer.salk.edu) (Heinz et al., 2010). For general analyses, we considered genes with reads over the noise threshold (20 reads). Significant expression in the γ neurons considered for genes with reads over a second noise threshold (50 reads) in at least two γ neuron. For normalization and statistics, we performed DEseq2 algorithm (Love et al., 2014) on our samples on R platform, which took into account batch effects. All p values presented for RNA-seq data are adjusted p values. Gene enrichment analysis was done using FlyMine (http://www.flymine.org/).

### Generation of CRISPR mediated mutant

For Dpr12 mutation two guide RNA were designed using the FlyCRISPR algorithm (http://flycrispr.molbio.wisc.edu/) and cloned into pCFD4 using Transfer-PCR (TPCR) (Unger et al., 2010, Melzer et al, under revision). The pCFD4-Dpr12 plasmid was injected into 86FB landing sites using ΦC31 integration (BestGene). Injected flies were crossed with nanos-Cas9 flies (Bloomington stock #54591). After two generations, single males were crossed with balancers and checked for deletion using specific primers. The Dpr12^Δ50-81^ allele is a 32bp deletion in the 5’ end of the transcript resulting in a premature stop after 37aa.

For tissue-specific CRISPR (tsCRISPR) Dpr12 and DIP-δ gRNA containing flies were crossed with UAS-Cas9.C or UAS-Cas9.P2 respectively, and the indicated Gal4s.

### Generation of transgenes and transgenic flies

To generate UAS-Dpr12 and UAS-DIP-δ transgenes, cDNA was cloned into the Gateway entry vector pDONR201. The Gateway entry vectors were then recombined into pDEST-UAS-IVS-Syn21-p10aw destination vector (Rabinovich et al., 2016) using LR recombinase (Invitrogen). UAS-Dpr12 and UAS-DIP-δ plasmids were injected into 86FB and attp40 landing sites, respectively, using ΦC31 integration (BestGene). For the generation of UAS-DIP-δ RNAi, 21 nucleotide sequence was selected using DSIR (http://biodev.extra.cea.fr/DSIR/DSIR.php). Off target results were eliminated by blast in NCBI. The RNAi hairpins were cloned into pVALIUM22 as described in https://fgr.hms.harvard.edu/cloning-and-sequencing. In brief, hairpins oligos, containing sense and anti-sense nucleotide with overhangs DNA fragment for *NheI* and *EcoRI* were synthesized (Sigma). 10 µl of sense and antisense strand oligos (10-20 µM each) were annealed into 80 µl annealing buffer (10 mM Tris-HCL, pH7.5, 0.1M NaCl, 1mM EDTA) by incubation at 95°C for 5 min. 6 µl of the annealed oligos were directly cloned in to pVALIUM22 vector which has been linearized by *NheI* and *EcoRI*. DIP-δ hairpins oligos (CAPS represent gene specific sequences): ctagcagtGACGATAAGAAACCTACAATAtagttatattcaagcataTTGTAGGTTTCTTATCGTC AGgcg aattcgcCTGACGATAAGAAACCTACAAtatgcttgaatataactaTATTGTAGGTTTCTTATCGT Cactg UAS-DIP-δ RNAi plasmid was injected into attp40 landing sites, using ΦC31 integration (BestGene).

### Generation of *DIP-δ*^*T2A-Gal4*^

*DIP-δ*^*T2A-Gal4*^ was generated as described in (Diao et al., 2015). In brief, flies carrying the MiMICMI08287 insertion were crossed with the flies bearing the triplet donor cassettes (Trojan Gal4 cassettes of the three reading frames). Males from this progeny carrying both components were crossed to females carrying germline transgenic sources of Cre and ΦC31. Adult progeny with all relevant components were crossed to UAS-GFP balanced on the 3^rd^ chromosome. Single males from this final cross were screened by fluorescence microscopy for Gal4 expression.

### Generation of QF2 driver

To generate the R71G10-QF2 driver, the QF2 sequence was amplified from pattB-DSCP_prom-QF7-hsp70_term (a gift from Chris potter) using the QF-F and QF-R primers, and cloned into pBPGUw plasmid, using the Gibson assembly kit (NEB) to create pBPGUw-QF2. Then the GMR71G10 entry vector (Alyagor et al., 2018)was recombined into pBPGUw-QF2 using LR recombinase (Invitrogen). Finally, the DSCP promotor region within the GMR71G10-QF2 was replaced with hsp70 promotor by RF cloning using the hspF and hspR primers. The GMR71G10-QF2^hsp70^ plasmid was injected into attp40 landing sites, using ΦC31 integration (BestGene).

QF-F TAAGCCAACTTTGAATCACAAGACGCATACCAAACGGTACATGCCACCCAAG QF-R TGAATAATTTTCTATTTGGCTTTAGTCGACGGTATCGATAATCACTGTTCGT HspF AAGTGGTGATAAACGGCCGGCCGAGCGCCGGAGTATAAATAGAG HspR AAGTGGTGATAAACGGCCGGCCGAGCGCCGGAGTATAAATAGAG

### Generation of MARCM clones

Due to Dpr12 centromeric chromosomal location we could not generate dpr12 mutant clones and instead expressed dpr12-RNAi within γ neuron clones. MB γ neuron MARCM clones were generated as described in (Lee and Luo, 1999). In brief, flies were heat shocked (hs) for 40-60 min at 37° C at 24hr after egg laying and examined at the indicated developmental time points.

To discover the mitotic window which results in adult PAM neurons, we used the MARCM technique to generate *DIP-δ*^*T2A-Gal4*^ clones by hs at different developmental times. Only hs at 0-24hr after egg laying resulted in clones containing PAM neurons, and therefore this hs regime was used in this study. Importantly, here mitotic recombination was performed using FRT40A and used to eliminate Gal80 expression but *DIP-δ* remained heterozygous as it is on another chromosome.

### Immunostaining and imaging

Brains were dissected in ringer solution, fixed using 4% paraformaldehyde (PFA) for 20 minutes at room temperature (RT) and washed with PB with 0.3% Triton-X (PBT, 3 immediate washes followed by 3 × 20 minute washes). Non-specific staining was blocked using 5% heat inactivated goat serum in PBT and then samples were subjected to primary antibodies (over-night, 4°C) and secondary antibodies (2 hours at RT) with PBT washes (3 quick washed followed by 3 × 20 minute washes). The brains were mounted on SlowFade (Invitrogen) and imaged using Zeiss LSM800 confocal microscope. Images were processed with ImageJ 1.51 (NIH). Individual neurons were traced manually through all focal planes using the Edge Detection Settings of the Analyze Paint Brush VVD selection tool (Takashi Kawase). Thresholding was individually adapted for each focal plane and neuronal structures and refined through the Analyze Erase tool whenever necessary.

For Brp staining brains were blocked in 2% bovine serum albumine (BSA) in PBT for 2 hr at RT, incubated with mouse anti-brp antibody for two days at 4°C then incubated with secondary antibody for an over-night at 4°C. Next, brains were transferred to a poly-L-lysine (Sigma Aldrich. # P1524-25MG) pre-treated cover glasses (22×22×1; Fisher Scientific. # 12-542B), fixed, dehydrated in ascending alcohol series (30%, 50%, 75%, 95% and 3 × 100%, 10 min each), incubated in Xylene 2 × 10 min and embedded in DPX (Electron Microscopy Sciences; # 180627-05) and incubated for at least 4 days. Confocal laser scanning microscopy was done using an Olympus microscope equipped with a Plan-Apochromat 20x objective. Taken images were analyzed using VVD Viewer (Takashi Kawase).

### Quantification and statistical analysis

In all cases, statistical significance was calculated as follows: *** represent a P-value lower than 0.001; ** represent a P-value lower than 0.01 and * represents a P-value lower than 0.05. Specific p-values and sample sizes are indicated in the relevant figure legends.

For quantification of regrowth (Figure 1, and 4), confocal Z-stacks were given to an independent lab member who blindly ranked the severity of the regrowth defects. For statistical analysis Kruskal-Wallis test was performed followed by a Wilcoxon–Mann– Whitney test post-hoc test.

## Supplemental Figures and Legends

**Figure S1.**
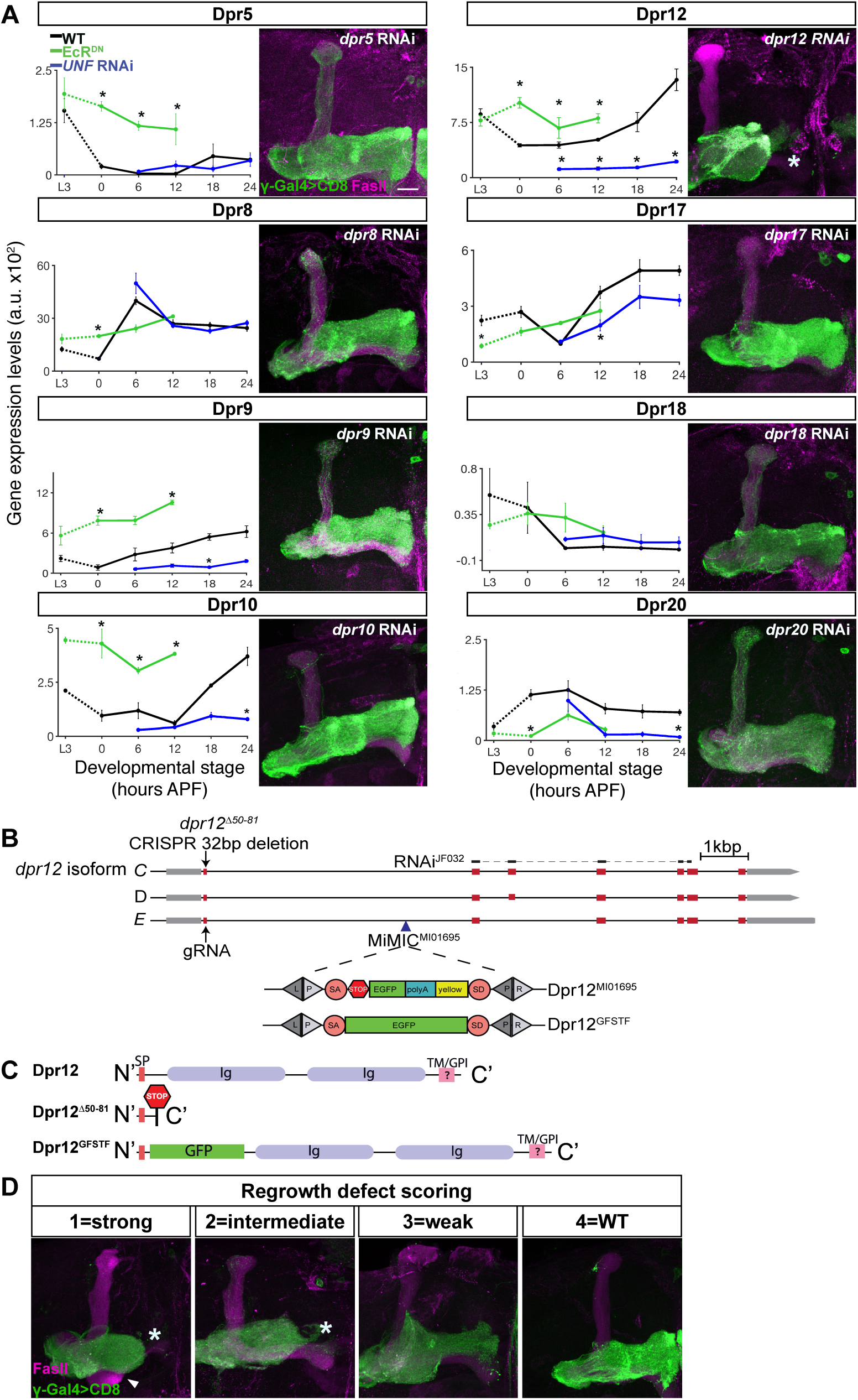
Knock down of *dpr12* results in regrowth phenotype and description of. ***Dpr12* alleles used in the study, related to Figure 1**. (A) Left: Graphs depicting the normalized RNA expression levels of selected Dprs in WT γ-KCs (black), and in γ-KCs expressing EcR^DN^ (green) or UNF-RNAi (blue). *p < 0.05; Error bars indicate SEM; units on the y axis are arbitrary. Right: Confocal z-projections of adult γ-KCs expressing RNAi transgenes as indicated labeled with membrane bound GFP (mCD8-GFP; CD8) driven by the γ specific Gal4 driver GMR71G10-Gal4 (γ-Gal4). (B) A schematic representation of the *Dpr12* locus showing introns (black line), coding and non-coding exons (red and gray, respectively). The location of *Dpr12* gRNA (arrow), *dpr12*^Δ*50-81*^ mutation (arrow), *dpr12* RNAi^JF03210^ (black lines connected with dashed lines) and MiMIC^MI01695^ (arrowhead) are indicated. SA and SD are splice acceptor and donor sites, respectively. Recombination mediated cassette exchange was used to transform Dpr12^MI01695^ into Dpr12^GFSTF^. (C) A schematic representation of Dpr12 protein variants. Signal peptide (SP), Immunoglobulin (Ig), transmembrane (TM) and GPI anchor (GPI). (D) Ranking of regrowth: Confocal z-projections of adult γ-KCs labeled with membrane bound GFP (mCD8-GFP; CD8) driven by the γ specific Gal4 driver GMR71G10-Gal4 (γ-Gal4). Representative images of the regrowth defect severity (1=strong, 2=intermediate, 3=weak, 4=WT) described in Figure 1O. Arrowhead demarcates short β lobe, which appear in approximately 40% of *dpr12* homozygous mutant brains, but not in *dpr12-RNAi* expressing brains. Asterisk demarcates the distal tip of the γ lobe. Green is CD8-GFP; magenta is FasII staining; Scale bar is 20µm.

**Figure S2.**
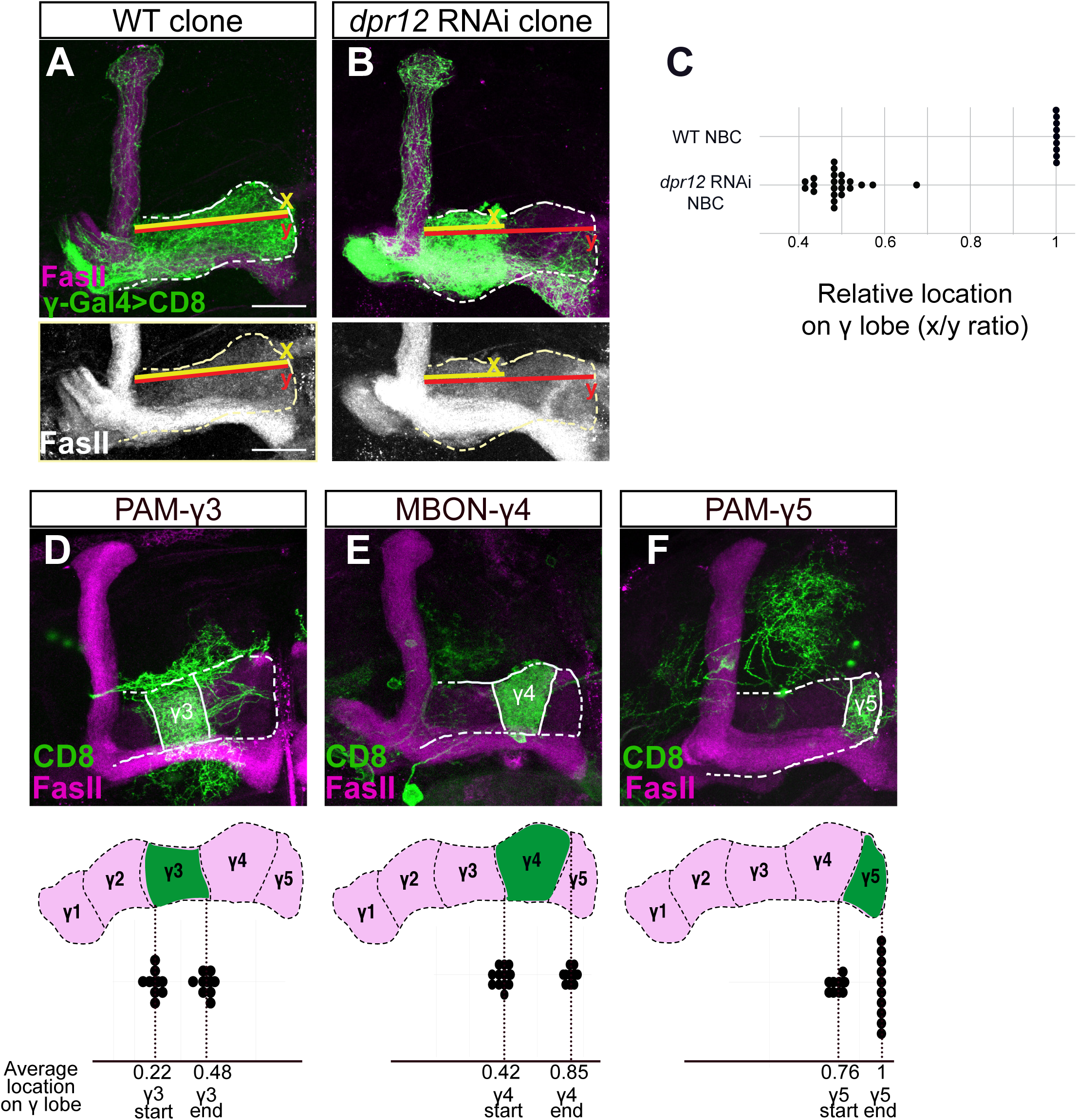
Measurements of γ-axon outgrowth, related to Figure 2. (A-B) Confocal z-projections of WT (A) and *dpr12* RNAi (B) MARCM neuroblast clones (NBC) labeled with membrane bound GFP (mCD8-GFP; CD8) driven by the γ specific Gal4 driver GMR71G10-Gal4 (γ-Gal4). y (red) represents the length of the entire γ-lobe as indicated; x (yellow) represents the extent of clonal γ-axon outgrowth. (C) Measurements of the x/y ration as depicted in A-B. While WT NBC always extend up to the end of the lobe, *dpr12* RNAi NBC stop at about midway (x/y ratio of 0.48 ± 0.05). (D-F) Top: Confocal z-projections of PAM-γ3 (D, MB441B), MBON-γ4 (E, R18H09), and PAM-γ5 (F, R48H11) Gal4s driving the expression of mCD8-GFP (CD8). Bottom: start and end of the indicated zone is superimposed on a schematic representation of the adult γ lobe compartments. (D) γ3 zone begins at x/y ratio of 0.22 ± 0.03 and ends at 0.48 ± 0.03. (E) γ4 zone begins at x/y ratio of 0.42 ± 0.04 and ends at 0.85 ± 0.03. (F) γ5 zone begins at x/y ratio of 0.76 ± 0.03 and ends at a mean ratio of 1. Green is CD8-GFP; magenta and white represent FasII; Scale bar is 20µm.

**Figure S3.**
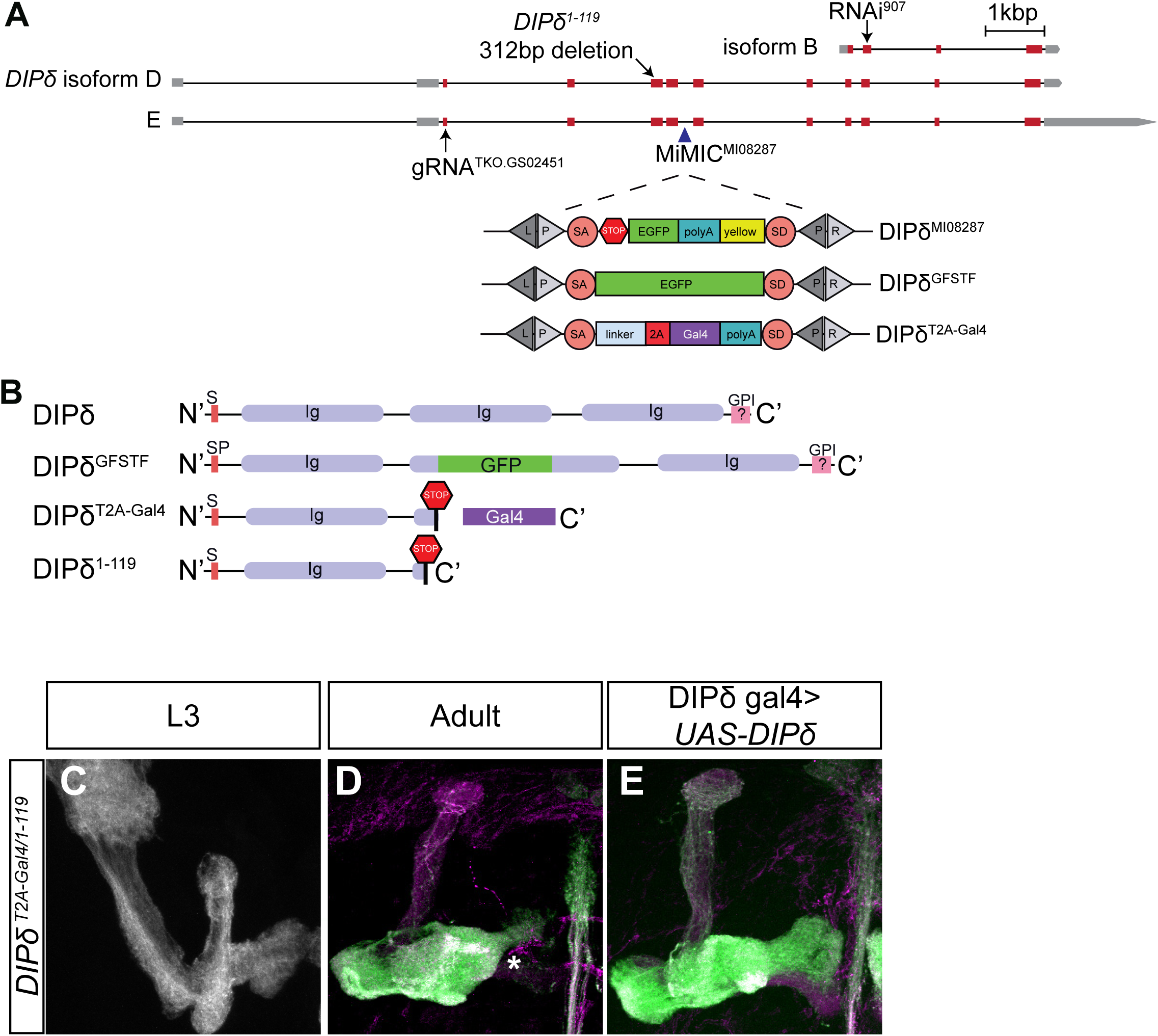
Description of *DIP-δ* alleles used in the study and additional *DIP-δ* loss of function phenotypes, related to Figure 4. (A) A schematic representation of the *DIP-δ* locus showing introns (black line), coding and non-coding exons (red and gray, respectively). The location of *DIP-δ* gRNA, *DIP-δ*^*1-119*^ mutation, *DIP-δ* RNAi and MiMIC^MI08287^ are indicated. SA and SD are splice acceptor and donor sites, respectively. Recombination mediated cassette exchange was used to transform DIP-δ^MI08287^ into DIP-δ^GFSTF^ and DIP-δ^T2A-Gal4^. (B) A schematic description of DIP-δ protein variants. Signal peptide (SP), Immunoglobulin (Ig), and GPI anchor (GPI). (C-E) Confocal z-projections of DIP-δ transheterozygotes (DIP-δ^T2A-Gal4/1-119^) at L3 (C; n=20/20) and adult (D; n=12/12, E; n=22/23), in which in which γ neurons were marked by expressing membrane bound tandem tomato (mtdT-HA) driven by the γ specific QF2 driver GMR71G10-QF2 (γ-QF2). In DIP-δ transheterozygotes, as in homozygous mutants (see Figure 4), γ neurons do not extend into the distal end of the lobe. Expression of DIP-δ in DIP-δ^+^ cells (E), rescues the growth defect present in DIP-δ^T2A-^ Gal4/1-119 brains. Green and white represent mtdT-HA; magenta represents FasII staining; Scale bar is 20µm.

**Figure S4.**
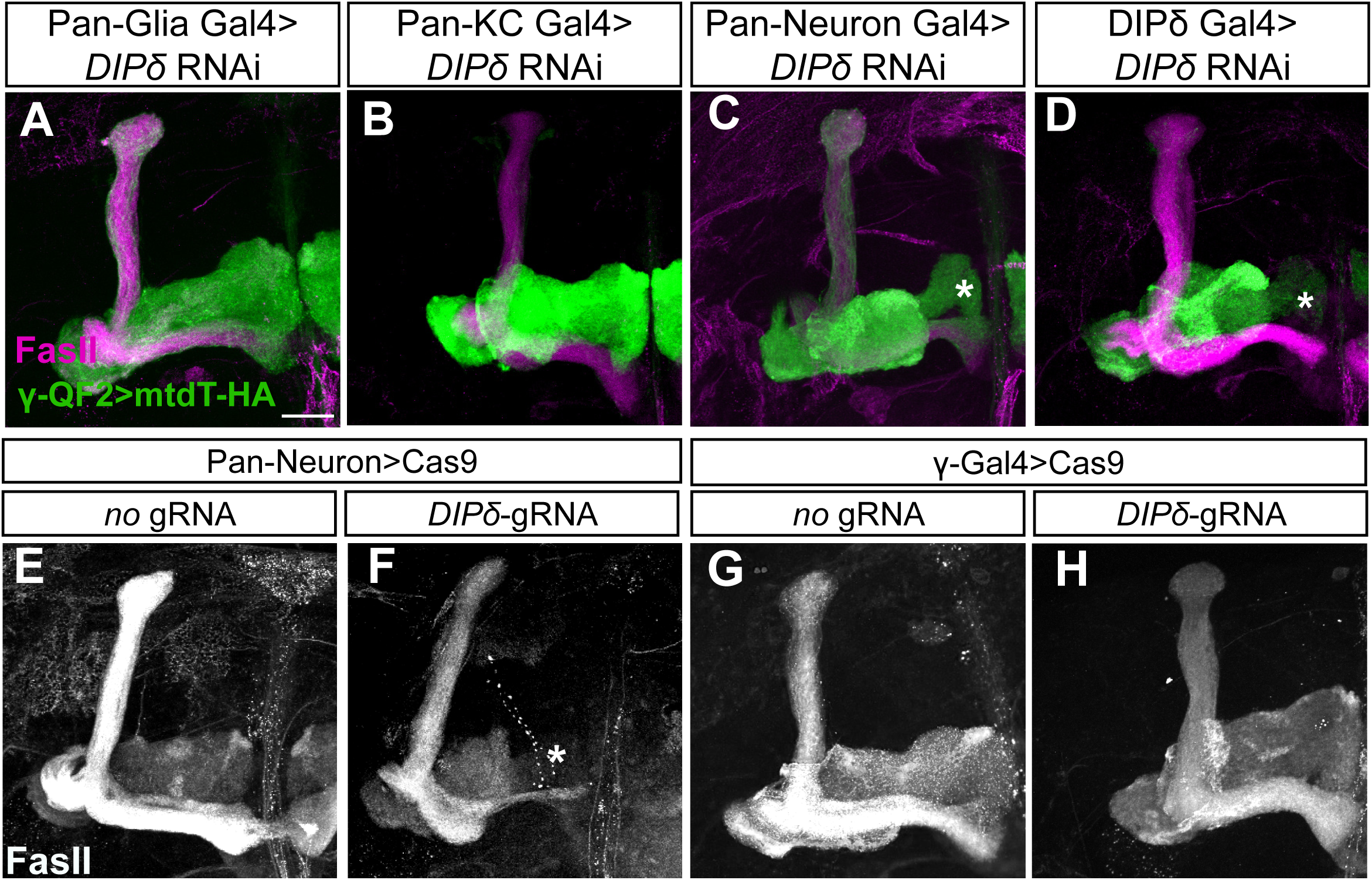
DIP-δ is non-cell-autonomously required for γ-axon regrowth, related to Figure 4. (A-D) Confocal z-projections of brains expressing *DIP-δ-*RNAi driven by the indicated Gal4. γ-KCs are labeled by membrane bound tandem tomato (mtdT-HA) driven by the γ specific QF2 driver GMR71G10-QF2 (γ-QF2). Expression of *DIP-δ*-RNAi in all glia (Repo-Gal4, L; n=28/28) or all KCs (OK107-Gal4, M; n=12/12) did not affect γ neuron regrowth. In contrast, expression of *DIP-δ*-RNAi in all postmitotic neurons (C155-Gal4, N; n=9/12) or DIP-δ expressing neurons (*DIP-δ*^*T2A-Gal4*^, O; n=21/22) induced a defect in γ4/5 innervation by γ-axons. (E-H) Confocal z-projections of brains expressing UAS-Cas9 alone (E,G) or together with *DIP-δ-*gRNA (F,H). *DIP-δ* knockout by tsCRISPR in all postmitotic neurons (F; n=22/24) resulted in a defect in γ4/5 innervation by γ-axons, while *DIP-δ* knockout by tsCRISPR in γ-KCs (H; n=28/28) did not affect γ-axon regrowth. Expression of Cas9 alone (E, n=10/10; G, n=14/14) did not affect γ-axon regrowth. Green is mtdT-HA; magenta and white represent FasII staining; Scale bar is 20µm.

**Figure S5.**
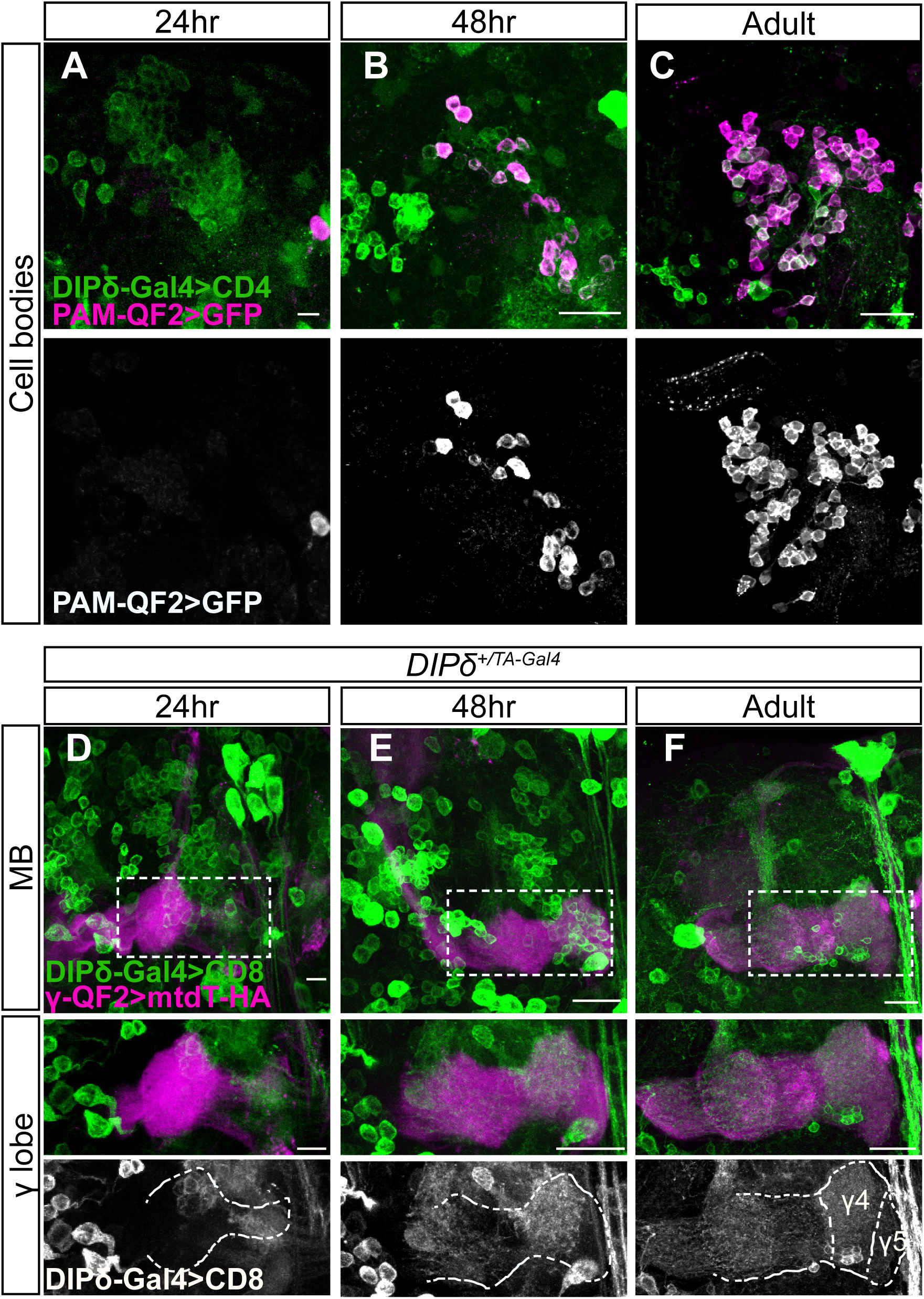
Characterization of PAM and DIP-δ Gal4s, related to Figure 5. (A-C) Confocal z-projections of the cell body region of the PAM cluster demonstrating the expression of membrane bound tandem tomato (CD4-tdT; CD4) driven by *DIP-δ*^*T2A-Gal4*^ (DIP-δ-Gal4) in addition to GFP driven by the PAM specific QF2 driver GMR58E02-QF2 (PAM-QF2). The PAM-QF2 driver is not expressed at 24hr APF (A, n=12), starts to be expressed at 48hr APF (B, n=10) and fully expressed and localized with DIP-δ-Gal4 in adult (C, n=14). (D-F) Confocal z-projections of heterozygous brains (DIP-δ^+/T2A-Gal4^) in which DIP-δ positive neurons were labeled by membrane bound GFP (mCD8-GFP; CD8) driven by DIP-δ-Gal4. γ-KCs were marked by expressing membrane bound tandem tomato (mtdT-HA) driven by the γ specific QF2 driver GMR71G10-QF2 (γ-QF2). Bottom: high magnification images, as demarcated by dashed box in top panels. Dashed outline demarcates γ-lobe as depicted by FasII staining. γ4/5 zones are indicated in adult. D, n=20; E, n=16; F, n=26. Scale bar is 20µm.

**Figure S6.**
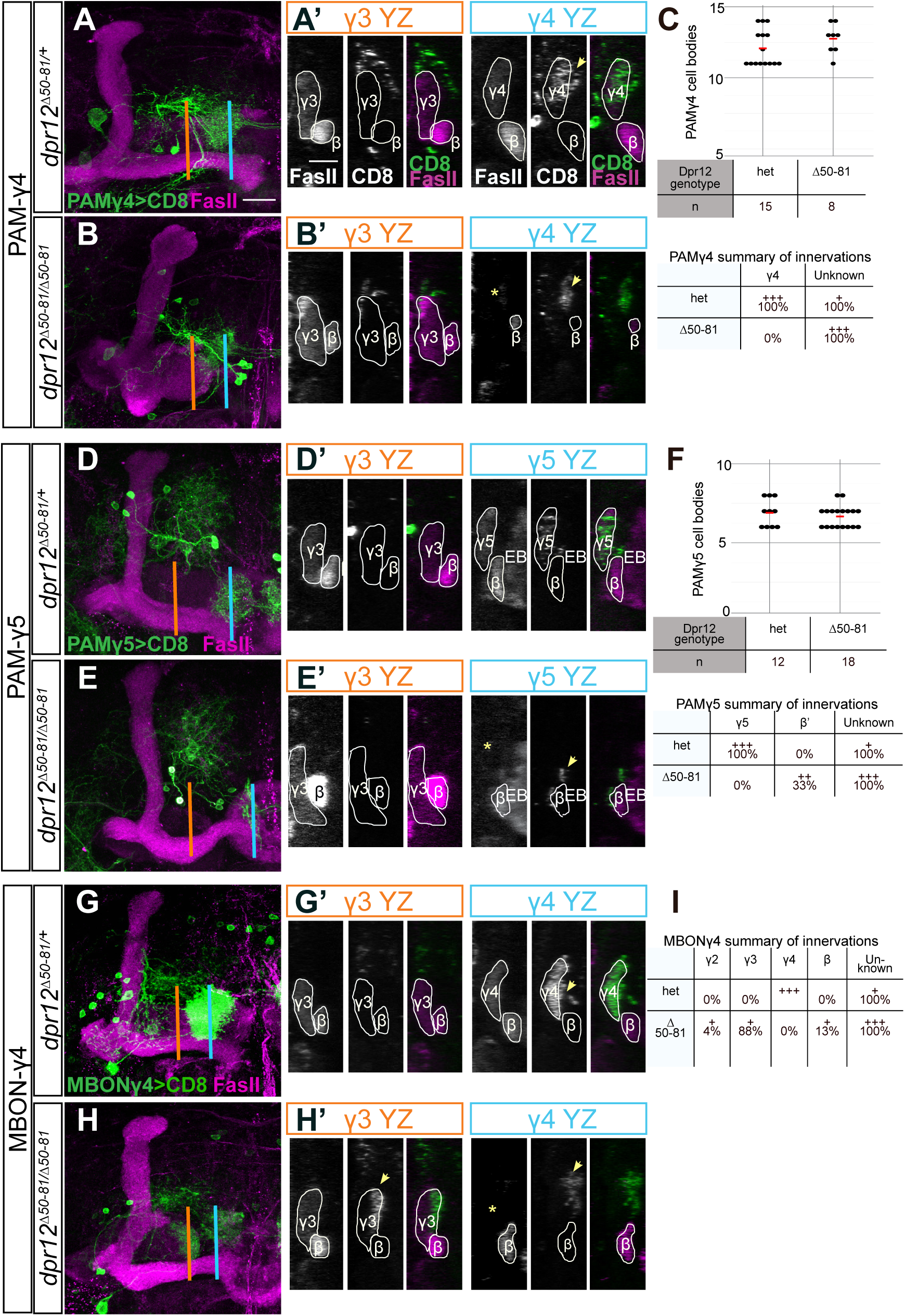
Phenotypic analysis of PAM and MBON innervation of the MB γ-lobe in Dpr12 mutant brains, related to Figure 7. (A-B, D-E, G-H) Left: Confocal z-projections of *dpr12* heterozygous or homozygous mutant brains (also shown in Figure 6) expressing membrane bound GFP (CD8) driven by: (A-B) R10G03-Gal4 is used to label PAM neurons innervating the γ4 compartment (PAM-γ4); (D-E) R48H11-Gal4 is used to label PAM neurons innervating the γ5 compartment (PAM-γ5); (G-H) R18H09-Gal4 is used to label MBON innervating the γ4 compartment (MBON-γ4). Right: YZ projections along the indicated lines in γ3 (orange) and γ4 or γ5 (blue) compartments. (C,F,I) Top: Cell body numbers of the indicated neurons in *dpr12* heterozygous and homozygous brains; Bottom: Summary of innervation destinations. Unknown means stereotypic projections to unidentifiable domains. Green is CD8-GFP, magenta is FasII, grayscale single channels are shown as indicated. Asterisks mark missing innervation and arrows mark innervations outside the γ lobe. Scale bar is 20µm in A-B, D-E, G-H and 10µm in A’-B’,D’-E’,G’-H’.

**Movie legends and Table descriptions**

**Movie S1: Tracing of control γ single cell clones**

Confocal reconstructions control MARCM γ single cell clones. Stacks were taken from Figure 2I, and used to trace single axons in Figure 2I’ which were included in the quantification in Figure 2N.

**Movie S2: Tracing of *dpr12-RNAi* γ single cell clones**

Confocal reconstructions MARCM γ single cell clones expressing *dpr12-RNAi*. Stacks were taken from Figure 2L, and used to trace single axons in Figure 2L’ which were included in the quantification in Figure 2N.

**Movie S3: Structure of γ-lobe compartments in WT brain**

Confocal reconstruction of a control brain (w^*^) stained with anti-Brp (the same brain that is presented in Figure 7G). The adult γ-lobe compartments (γ1-γ5) are marked with yellow, g green, cyan, blue and light blue. The adult αβ and α’β’ neurons are marked with magenta and red respectively. Scale bar is 20µm.

**Movie S4: Structure of γ-lobe compartments in *dpr12* mutant brain**

Confocal reconstruction of *dpr12* homozygous mutant brain stained with anti-Brp (as presented at Figure 7H). The adult γ-lobe compartments (γ1-γ4) are marked with yellow, green, cyan, and blue. Note that the γ5 and most of the γ4 compartments are missing. The adult αβ and α’β’ neurons are marked with magenta and red respectively. Scale bar is 20µm.

**Table S1: Normalized RNAseq data**

Normalized and averaged RNAseq profiling of WT MB γ neurons during development (taken from ref 14), compared to expression profiles of MB γ neurons expressing UNF-RNAi.

**Table S2: Expression analysis of the Dpr/DIP families**

Normalized and averaged RNAseq values of Dpr and DIP transcripts in WT MB γ neurons during development (taken from ref 14), compared to expression profiles of MB γ neurons expressing UNF-RNAi.

## References

Alyagor, I., Berkun, V., Keren-Shaul, H., Marmor-Kollet, N., David, E., Mayseless, O., Issman-Zecharya, N., Amit, I., and Schuldiner, O. (2018). Combining Developmental and Perturbation-Seq Uncovers Transcriptional Modules Orchestrating Neuronal Remodeling. Developmental cell 47, 38–52 e36.

Ashley, J., Sorrentino, V., Lobb-Rabe, M., Nagarkar-Jaiswal, S., Tan, L., Xu, S., Xiao, Q., Zinn, K., and Carrillo, R.A. (2019). Transsynaptic interactions between IgSF proteins DIP-alpha and Dpr10 are required for motor neuron targeting specificity. eLife 8.

Aso, Y., Hattori, D., Yu, Y., Johnston, R.M., Iyer, N.A., Ngo, T.T., Dionne, H., Abbott, L.F., Axel, R., Tanimoto, H., et al. (2014a). The neuronal architecture of the mushroom body provides a logic for associative learning. eLife 3, e04577.

Aso, Y., Sitaraman, D., Ichinose, T., Kaun, K.R., Vogt, K., Belliart-Guerin, G., Placais, P.Y., Robie, A.A., Yamagata, N., Schnaitmann, C., et al. (2014b). Mushroom body output neurons encode valence and guide memory-based action selection in Drosophila. eLife 3, e04580.

Bilz, F., Geurten, B.R.H., Hancock, C.E., Widmann, A., and Fiala, A. (2020). Visualization of a Distributed Synaptic Memory Code in the Drosophila Brain. Neuron.

Carrillo, R.A., Ozkan, E., Menon, K.P., Nagarkar-Jaiswal, S., Lee, P.T., Jeon, M., Birnbaum, M.E., Bellen, H.J., Garcia, K.C., and Zinn, K. (2015). Control of Synaptic Connectivity by a Network of Drosophila IgSF Cell Surface Proteins. Cell 163, 1770–1782.

Cognigni, P., Felsenberg, J., and Waddell, S. (2018). Do the right thing: neural network mechanisms of memory formation, expression and update in Drosophila. Current opinion in neurobiology 49, 51–58.

Cohn, R., Morantte, I., and Ruta, V. (2015). Coordinated and Compartmentalized Neuromodulation Shapes Sensory Processing in Drosophila. Cell 163, 1742–1755.

Cosmanescu, F., Katsamba, P.S., Sergeeva, A.P., Ahlsen, G., Patel, S.D., Brewer, J.J., Tan, L., Xu, S., Xiao, Q., Nagarkar-Jaiswal, S., et al. (2018). Neuron-Subtype-Specific Expression, Interaction Affinities, and Specificity Determinants of DIP/Dpr Cell Recognition Proteins. Neuron 100, 1385–1400 e1386.

Crittenden, J.R., Skoulakis, E.M., Han, K.A., Kalderon, D., and Davis, R.L. (1998). Tripartite mushroom body architecture revealed by antigenic markers. Learning & memory 5, 38–51.

Croset, V., Treiber, C.D., and Waddell, S. (2018). Cellular diversity in the Drosophila midbrain revealed by single-cell transcriptomics. eLife 7.

Fiala, A. (2007). Olfaction and olfactory learning in Drosophila: recent progress. Current opinion in neurobiology 17, 720–726.

Gerber, B., Scherer, S., Neuser, K., Michels, B., Hendel, T., Stocker, R.F., and Heisenberg, M. (2004). Visual learning in individually assayed Drosophila larvae. J Exp Biol 207, 179–188.

Heisenberg, M. (2003). Mushroom body memoir: from maps to models. Nature reviews Neuroscience 4, 266–275.

Lee, T., Lee, A., and Luo, L. (1999). Development of the Drosophila mushroom bodies: sequential generation of three distinct types of neurons from a neuroblast. Development 126, 4065–4076.

Lee, T., Marticke, S., Sung, C., Robinow, S., and Luo, L. (2000). Cell-autonomous requirement of the USP/EcR-B ecdysone receptor for mushroom body neuronal remodeling in Drosophila. Neuron 28, 807–818.

Meltzer, H., Marom, E., Alyagor, I., Mayseless, O., Berkun, V., Segal-Gilboa, N., Unger, T., Luginbuhl, D., and Schuldiner, O. (2019). Tissue-specific (ts)CRISPR as an efficient strategy for in vivo screening in Drosophila. Nature communications 10, 2113.

Modi, M.N., Shuai, Y., and Turner, G.C. (2020). The Drosophila Mushroom Body: From Architecture to Algorithm in a Learning Circuit. Annual review of neuroscience.

Nagarkar-Jaiswal, S., Lee, P.T., Campbell, M.E., Chen, K., Anguiano-Zarate, S., Gutierrez, M.C., Busby, T., Lin, W.W., He, Y., Schulze, K.L., et al. (2015). A library of MiMICs allows tagging of genes and reversible, spatial and temporal knockdown of proteins in Drosophila. eLife 4.

Owald, D., and Waddell, S. (2015). Olfactory learning skews mushroom body output pathways to steer behavioral choice in Drosophila. Current opinion in neurobiology 35, 178–184.

Ozkan, E., Carrillo, R.A., Eastman, C.L., Weiszmann, R., Waghray, D., Johnson, K.G., Zinn, K., Celniker, S.E., and Garcia, K.C. (2013). An extracellular interactome of immunoglobulin and LRR proteins reveals receptor-ligand networks. Cell 154, 228–239.

Port, F., and Bullock, S.L. (2016). Augmenting CRISPR applications in Drosophila with tRNA-flanked sgRNAs. Nature methods 13, 852–854.

Port, F., Strein, C., Stricker, M., Rauscher, B., Heigwer, F., Zhou, J., Beyersdorffer, C., Frei, J., Hess, A., Kern, K., et al. (2020). A large-scale resource for tissue-specific CRISPR mutagenesis in Drosophila. eLife 9.

Rabinovich, D., Yaniv, S.P., Alyagor, I., and Schuldiner, O. (2016). Nitric Oxide as a Switching Mechanism between Axon Degeneration and Regrowth during Developmental Remodeling. Cell 164, 170–182.

Rohwedder, A., Wenz, N.L., Stehle, B., Huser, A., Yamagata, N., Zlatic, M., Truman, J.W., Tanimoto, H., Saumweber, T., Gerber, B., et al. (2016). Four Individually Identified Paired Dopamine Neurons Signal Reward in Larval Drosophila. Current biology : CB 26, 661–669.

Sanz, R., Ferraro, G.B., and Fournier, A.E. (2015). IgLON cell adhesion molecules are shed from the cell surface of cortical neurons to promote neuronal growth. The Journal of biological chemistry 290, 4330–4342.

Saumweber, T., Rohwedder, A., Schleyer, M., Eichler, K., Chen, Y.C., Aso, Y., Cardona, A., Eschbach, C., Kobler, O., Voigt, A., et al. (2018). Functional architecture of reward learning in mushroom body extrinsic neurons of larval Drosophila. Nature communications 9, 1104.

Shuai, Y., Hirokawa, A., Ai, Y., Zhang, M., Li, W., and Zhong, Y. (2015). Dissecting neural pathways for forgetting in Drosophila olfactory aversive memory. Proceedings of the National Academy of Sciences of the United States of America 112, E6663–6672.

Tan, L., Zhang, K.X., Pecot, M.Y., Nagarkar-Jaiswal, S., Lee, P.T., Takemura, S.Y., McEwen, J.M., Nern, A., Xu, S., Tadros, W., et al. (2015). Ig Superfamily Ligand and Receptor Pairs Expressed in Synaptic Partners in Drosophila. Cell 163, 1756–1769.

Tanaka, N.K., Tanimoto, H., and Ito, K. (2008). Neuronal assemblies of the Drosophila mushroom body. J Comp Neurol 508, 711–755.

Venkatasubramanian, L., Guo, Z., Xu, S., Tan, L., Xiao, Q., Nagarkar-Jaiswal, S., and Mann, R.S. (2019). Stereotyped terminal axon branching of leg motor neurons mediated by IgSF proteins DIP-alpha and Dpr10. eLife 8.

Xu, S., Xiao, Q., Cosmanescu, F., Sergeeva, A.P., Yoo, J., Lin, Y., Katsamba, P.S., Ahlsen, G., Kaufman, J., Linaval, N.T., et al. (2018). Interactions between the Ig-Superfamily Proteins DIP-alpha and Dpr6/10 Regulate Assembly of Neural Circuits. Neuron 100, 1369–1384 e1366.

Yaniv, S.P., Issman-Zecharya, N., Oren-Suissa, M., Podbilewicz, B., and Schuldiner, O. (2012). Axon regrowth during development and regeneration following injury share molecular mechanisms. Current biology : CB 22, 1774–1782.

Yaniv, S.P., Meltzer, H., Alyagor, I., and Schuldiner, O. (2020). Developmental axon regrowth and primary neuron sprouting utilize distinct actin elongation factors. The Journal of cell biology 219.

Zinn, K., and Ozkan, E. (2017). Neural immunoglobulin superfamily interaction networks. Current opinion in neurobiology 45, 99–105.

## Supplemental References

Alyagor, I., Berkun, V., Keren-Shaul, H., Marmor-Kollet, N., David, E., Mayseless, O., Issman-Zecharya, N., Amit, I., and Schuldiner, O. (2018). Combining Developmental and Perturbation-Seq Uncovers Transcriptional Modules Orchestrating Neuronal Remodeling. Dev Cell 47, 38–52 e36.

Diao, F., Ironfield, H., Luan, H., Diao, F., Shropshire, W.C., Ewer, J., Marr, E., Potter, C.J., Landgraf, M., and White, B.H. (2015). Plug-and-play genetic access to drosophila cell types using exchangeable exon cassettes. Cell Rep 10, 1410–1421.

Gratz, S.J., Ukken, F.P., Rubinstein, C.D., Thiede, G., Donohue, L.K., Cummings, A.M., and O’Connor-Giles, K.M. (2014). Highly specific and efficient CRISPR/Cas9-catalyzed homology-directed repair in Drosophila. Genetics 196, 961–971.

Heinz, S., Benner, C., Spann, N., Bertolino, E., Lin, Y.C., Laslo, P., Cheng, J.X., Murre, C., Singh, H., and Glass, C.K. (2010). Simple combinations of lineage-determining transcription factors prime cis-regulatory elements required for macrophage and B cell identities. Mol Cell 38, 576–589.

Kim, D., Langmead, B., and Salzberg, S.L. (2015). HISAT: a fast spliced aligner with low memory requirements. Nat Methods 12, 357–360.

Lee, T., and Luo, L. (1999). Mosaic analysis with a repressible cell marker for studies of gene function in neuronal morphogenesis. Neuron 22, 451–461.

Love, M.I., Huber, W., and Anders, S. (2014). Moderated estimation of fold change and dispersion for RNA-seq data with DESeq2. Genome Biol 15, 550.

Port, F., Chen, H.M., Lee, T., and Bullock, S.L. (2014). Optimized CRISPR/Cas tools for efficient germline and somatic genome engineering in Drosophila. Proc Natl Acad Sci U S A 111, E2967–2976.

Vert, J.P., Foveau, N., Lajaunie, C., and Vandenbrouck, Y. (2006). An accurate and interpretable model for siRNA efficacy prediction. BMC Bioinformatics 7, 520.

